# Efficient Generation of Protein Pockets with PocketGen

**DOI:** 10.1101/2024.02.25.581968

**Authors:** Zaixi Zhang, Wan Xiang Shen, Qi Liu, Marinka Zitnik

## Abstract

Designing protein-binding proteins is critical for drug discovery. However, the AI-based design of such proteins is challenging due to the complexity of ligand-protein interactions, the flexibility of ligand molecules and amino acid side chains, and sequence-structure dependencies. We introduce PocketGen, a deep generative model that simultaneously produces both the residue sequence and atomic structure of the protein regions where ligand interactions occur. PocketGen ensures consistency between sequence and structure by using a graph transformer for structural encoding and a sequence refinement module based on a protein language model. The bilevel graph transformer captures interactions at multiple scales, including atom, residue, and ligand levels. To enhance sequence refinement, PocketGen integrates a structural adapter into the protein language model, ensuring that structure-based predictions align with sequence-based predictions. PocketGen can generate high-fidelity protein pockets with superior binding affinity and structural validity. It operates ten times faster than physics-based methods and achieves a 95% success rate, defined as the percentage of generated pockets with higher binding affinity than reference pockets. Additionally, it attains an amino acid recovery rate exceeding 64%.

## Introduction

Modulating protein functions involves modeling the interactions between proteins and small molecule ligands [1, 2, 3, 4]. These interactions are central to biological processes such as enzymatic catalysis, signal transduction, and cellular regulation. Binding small molecules to specific protein sites can induce conformational changes, modulate protein activity, and alter existing or produce new functional properties. This mechanism is invaluable for studying protein functions and designing proteins with tailored small molecule-binding properties. Applications range from engineering enzymes to catalyze reactions in the absence of natural catalysts [5, 6, 7, 8] to creating biosensors for detecting environmental compounds. Such biosensors are critical for environmental monitoring, clinical diagnostics, pathogen detection, drug delivery systems, and food industry applications [9, 10, 11, 12]. Typically, designs involve modifying existing ligand-binding pockets to enable more specific interactions with target ligands [13, 14, 15]. Nevertheless, challenges persist in computationally generating high-validity ligand-binding protein pockets due to the complexity of ligand-protein interactions, the flexibility of ligands and amino acid side chains, and the dependencies between sequence and structure [3, 15, 16].

Methods for pocket design have traditionally relied on physics-based modeling or template matching [10, 11, 13, 17, 18]. For example, PocketOptimizer [18, 19, 20] uses a pipeline that predicts mutations in protein pockets to enhance binding affinity, based on physics-based energy functions and search algorithms. Starting with a bound protein-ligand complex, PocketOptimizer explores possible side chain structures and residue types, evaluating these mutations with energy functions and ranking them using integer linear programming techniques. Another widely used approach involves template matching and enumeration methods [11, 13, 14, 17, 21]. For instance, Polizzi et al. [13] use a two-step strategy for pocket design. First, they identify and assemble disconnected protein motifs (van der Mer (vdM) structural units) around the target molecule to form protein-ligand hydrogen bonds. Then, they graft these residues onto a protein scaffold and select the optimal protein-ligand pairs using scoring functions. This template-matching strategy enabled the *de novo* design of proteins binding the drug apixaban [22]. However, physics-based and template-matching methods can be time-consuming, often requiring several hours to design a single protein pocket. Furthermore, the focus on specific fold types, such as four-helix bundles [13] or NTF2 folds [14], can limit the broader applicability of these methods.

Recent advances in protein pocket design have been propelled by deep learning-based approaches [3, 8, 16, 23, 24, 25]. For instance, RFDiffusion [26] leverages denoising diffusion probabilistic models [27] alongside RoseTTAFold [28] for *de novo* protein structure generation. Although it can design pockets for specific ligands, RFDiffusion lacks precision in modeling protein-ligand interactions due to its auxiliary guiding potentials. To address this limitation, RFdiffusion All-Atom (RFdiffusionAA) [16] extends the approach by enabling direct generation of binding proteins around small molecules through iterative denoising. This is achieved through architectural modifications that simultaneously consider both protein structures and ligand molecules. However, in both RFDiffusion and RFdiffusionAA, residue sequences are derived in post-processing using ProteinMPNN [29] or LigandMPNN [30], which can result in inconsistencies between the sequence and structure modalities. In contrast, FAIR [24] simultaneously designs the atomic pocket structure and the corresponding sequence using a two-stage refinement approach. FAIR employs a coarse-to-fine method, initially refining the backbone protein structure and subsequently refining the atomic structure, including the side chains. This iterative process continues until convergence is reached. However, the gap between these two refinement stages can introduce instability and limit performance, underscoring the need for an end-to-end generative approach to pocket design. Related research has explored the co-design of sequence and structure in complementarity determining regions (CDRs) of antibodies [31, 32, 33, 34, 35]. While these methods are effective for antibody design, they encounter difficulties when applied to pocket designs conditioned on target ligand molecules.

Hybrid approaches that combine deep learning models with traditional methods are also being actively explored [3, 8]. For example, Yeh et al. [8] developed a novel Luciferase by integrating protein hallucination [36], trRosetta structure prediction neural network [37], hydrogen bonding networks, and RifDock [38]. This combination generated a range of idealized protein structures with diverse pocket shapes for subsequent filtering. While successful, this approach applies only to specific protein scaffolds and substrates and lacks a generalized solution. Similarly, Lee et al. [3] merge deep learning with physics-based methods to design proteins featuring diverse and customizable pocket geometries. Their method utilizes backbone generation via trRosetta hallucination, sequence design through ProteinMPNN [29] and LigandMPNN [30], and filtering with AlphaFold [39]. Despite the advances made, pocket generation models continue to face challenges, such as achieving sequence-structure consistency and accurately modeling complex protein-ligand interactions.

Here, we introduce PocketGen, a deep generative method designed for efficient generation of protein pockets. PocketGen employs a co-design scheme (Figure 1a), where the model simultaneously predicts both the sequence and structure of the protein pocket based on the ligand molecule and the surrounding protein scaffold (excluding the pocket itself). The architecture of PocketGen is composed of two key modules: the bilevel graph transformer (Figure 1b) and the sequence refinement module (Figure 1c). PocketGen represents the protein-ligand complex as a geometric graph of blocks, enabling it to manage the variable atom counts across different residues and ligands. Initially, pocket residues are assigned the maximum possible number of atoms (14 atoms) to accommodate variability, and after generation, these atoms are mapped back to specific residue types.

**Figure 1.**
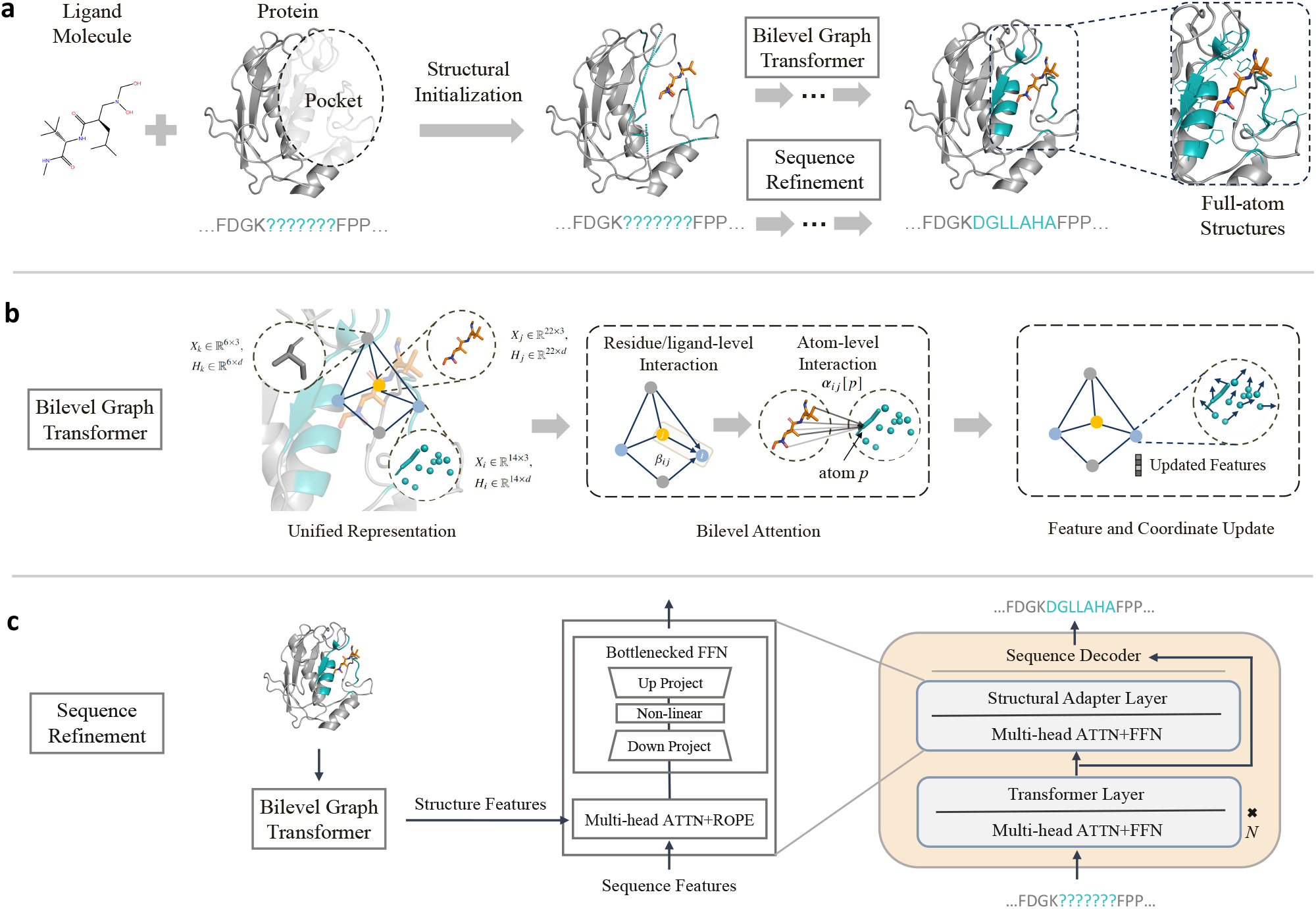
Overview of PocketGen generative model for the design of full-atom ligand-binding protein pockets. **a**, Conditioned on the binding ligand molecule and the rest part of the protein except the pocket region (i.e., scaffold), PocketGen aims to generate the full atom pocket structure (backbone and sidechain atoms) and the residue type sequence with iterative equivariant refinement. The ligand structure is also adjusted during the protein pocket refinement. **b**, Bilevel graph transformer is leveraged in PocketGen for all-atom structural encoding and update. The bilevel level attention captures both the residue/ligand and atom-level interactions. Both the protein pocket structure and the ligand molecule structure are updated in the refinement. **c**, Sequence refinement module adds lightweight structural adapter layers into pLMs for sequence prediction. Only the adapter’s parameters are fine-tuned during training, and the other layers are fixed. In the adapter, the cross-attention between sequence and structure features is performed to achieve sequence-structure consistency.

The graph transformer module uses a bilevel attention mechanism to capture interactions at multiple granularities—both at the atom and residue/ligand levels—and across various aspects, including intra-protein and protein-ligand interactions. To account for the redesigned pocket’s influence on the ligand, the ligand structure is updated during the refinement process to reflect potential changes in binding pose. To ensure consistency between the protein sequence and structure domains and to incorporate evolutionary information encoded in protein language models (pLMs) [40, 41], PocketGen integrates a structural adapter into the sequence update process. This adapter enables cross-attention between sequence and structure features, ensuring sequence-structure alignment. Only the adapter is fine-tuned during training, while the remaining layers of the protein language model remain unchanged. PocketGen outperforms existing methods for protein pocket generation across two popular benchmarks. It achieves an average amino acid recovery rate of 63.40% and a Vina score of -9.655 for top-1 ranked generated protein pockets on the CrossDocked dataset. Comprehensive analyses show that PocketGen can generate diverse, high-affinity protein pockets for functional molecules, highlighting its efficacy and potential for designing small-molecule binders and enzymes.

## Results

### Benchmarking generated protein pockets

We benchmark PocketGen on two datasets. The **CrossDocked** dataset [42] consists of protein-molecule pairs generated through cross-docking and is divided into training, validation, and test sets based on a 30% sequence identity threshold. The **Binding MOAD** dataset [43] contains experimentally determined protein-ligand complexes, which are split into training, validation, and test sets according to the proteins’ enzyme commission numbers [44]. In line with intermolecular distance scales relevant to protein-ligand interactions [45], our default experimental setup includes all residues with atoms within 3.5 Å of any ligand-binding atoms, averaging about eight residues per pocket. We also explore PocketGen’s ability to design larger pockets with a radius of 5.5 Å, incorporating more residues (Figure 3c).

We use three groups of metrics to evaluate the quality of protein pockets generated by PocketGen. First, we assess the affinity between the generated pocket and the target ligand molecule using the AutoDock **Vina score** [46], **MM-GBSA** [47], and **min-in-place GlideSP score** [48]. Second, we evaluate the structural validity of the generated pockets using **scRMSD, scTM**, and **pLDDT**. The amino acid sequence for the protein pocket structure is derived using ProteinMPNN [29], and the pocket structure is predicted using ESMFold [49] or AlphaFold2 [39]. The **scRMSD** is calculated as the self-consistency root mean squared deviation between the generated structure’s backbone atoms and the predicted structure. Following an established strategy [50, 51], eight sequences are predicted for each generated protein structure, and the sequence with the lowest scRMSD is used for reporting. Similarly, **scTM**, the self-consistency template modeling score, is calculated by comparing the TM-score [52] between the predicted and generated structures. Scores range from 0 to 1, with higher values indicating greater designability. We also report the Δ**scTM score** to assess whether the generated pocket improves or degrades the scTM score of the initial protein. The **pLDDT score** [39] reflects the confidence in structural predictions on a scale from 0 to 100, with higher scores indicating greater confidence. The average pLDDT score across pocket residues is reported. A generated protein pocket is defined as **designable** if the overall structure’s scRMSD is less than 2 Å and the pocket’s scRMSD is less than 1 Å [26, 53, 54]. Table S1 presents the percentage of designable generated pockets, and Supplementary Figure S1 describes how these metrics are calculated. Finally, we report the **amino acid recovery (AAR)** as the percentage of correctly predicted pocket residue types, which reflects the accuracy of the designed sequence. A higher AAR indicates better modeling of sequence-structure dependencies.

We compare PocketGen against six methods, including deep learning-based approaches such as RFDiffusion [26], RFDiffusionAA (RFAA) [16], FAIR [24], and dyMEAN [25], as well as a template-matching method, DEPACT [17], and a physics-based modeling method, PocketOpt [18] (Methods). In Figure 2 and Table S1, PocketGen and the other methods are tasked with generating 100 sequences and structures for each protein-ligand complex in the test sets of the CrossDocked and Binding MOAD datasets. PocketOpt is excluded from this comparison due to its focus on mutating existing pockets for optimization, making it too time-consuming to generate many protein pockets. Table S1 presents the mean and standard deviation of results across three independent runs with different random seeds. In Figure 2, we apply bootstrapping to the generation results, illustrating the distributions to demonstrate the sensitivity of the results to the dataset composition [55]. As shown in Table S1 and Figure 2, PocketGen outperforms all baselines, including RFDiffusion and RFDiffusionAA (RFAA), in terms of designability (by 3% and 2% on CrossDocked, respectively) and Vina scores (by 0.199 and 0.123 on CrossDocked, respectively). This performance indicates PocketGen’s effectiveness in generating structurally valid pockets with high binding affinities, a result attributed to PocketGen’s ability to capture interactions at multiple granularities—both atom-level and residue/ligand-level—and across various aspects, including intra-protein and protein-ligand interactions.

**Figure 2.**
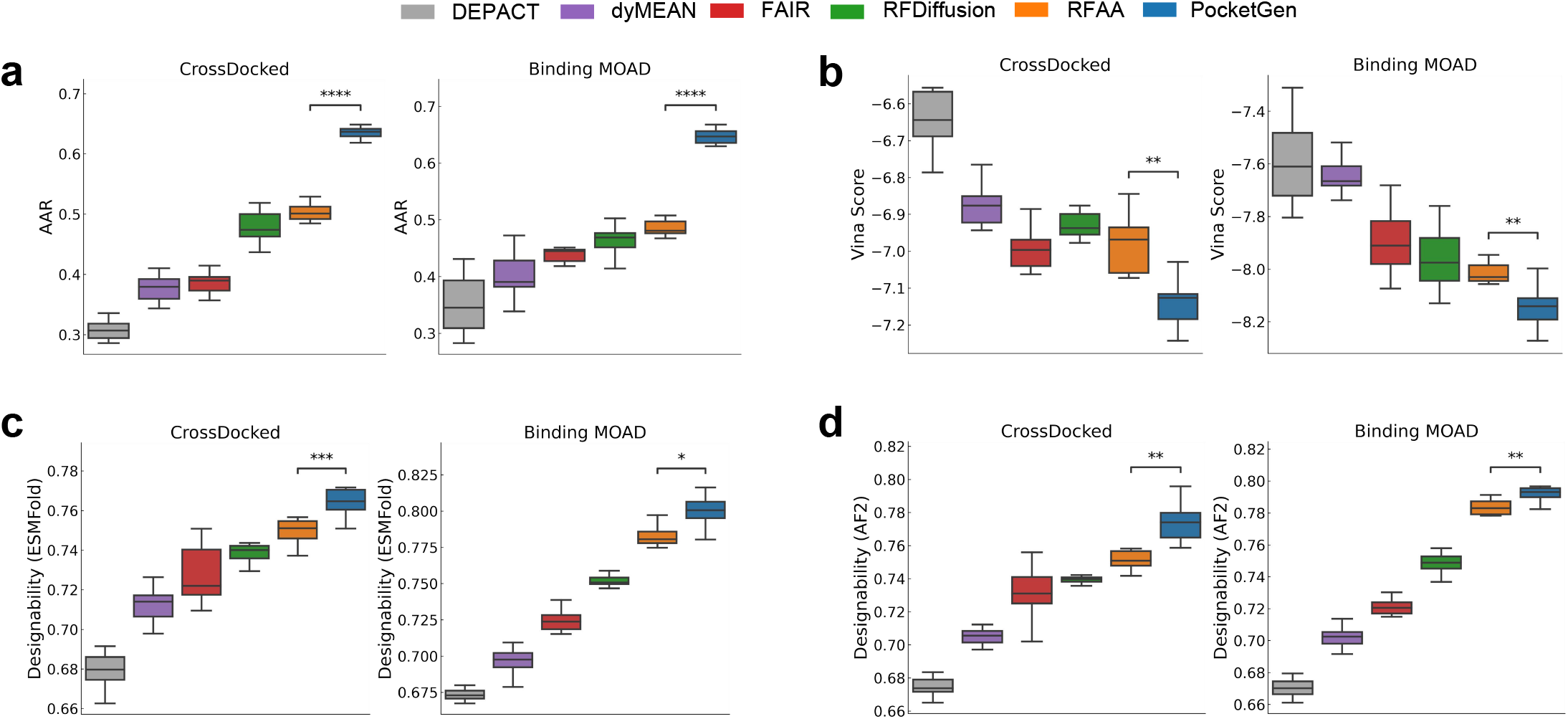
Benchmarking PocketGen on CrossDocked and Binding MOAD datasets. Shown are **a**, amino acid recovery rates (AAR) (p values 3.8e-8 and 1.5e-10), **b**, Vina score performance (p values 6.1e-3 and 6.7e-3), **c**, Designability scores using ESMFold structure prediction method (p values 6.0e-4 and 2.5e-2), and **d**, Designability scores using AF2 structure prediction method (p values 4.4e-3 and 4.4e-3). Uncertainty is quantified via bootstrapping, two-sided Kolmogorov-Smirnov test is used to compare PocketGen to the best-performing existing model (RFAA). P-value annotation legend: *: *p* ∈ [0, 01, 0.05], **: *p* ∈ [0.001, 0.01], ***: *p* ∈ [0.0001, 0.001], ****: *p* ≤ 0.0001. The sample size in the plots are 10 for each model. In all the box plots, the minimum is the smallest value within the data set, marked at the end of the lower whisker. The first quartile (Q1), or 25th percentile, forms the lower edge of the box. The median (50th percentile) is represented by a line inside the box, indicating the midpoint of the data. The third quartile (Q3), or 75th percentile, forms the upper edge of the box. The maximum is the largest value within the data set, marked at the end of the upper whisker. The whiskers extend to the smallest and largest values within 1.5 times the interquartile range (IQR).

PocketGen significantly outperforms the best-performing alternative method, RFDiffusionAA, with an average improvement of 13.95% in amino acid recovery rate (AAR), largely due to including a protein language model that captures evolutionary sequence information. In contrast, RFDiffusion and RFDiffusionAA rely on post-processing to determine amino acid types, which can lead to inconsistencies between sequence and structure and lower performance in AAR. In protein engineering, the common practice is to mutate several key residues to optimize properties while keeping most residues unchanged to preserve protein folding stability [56, 57]. The high AAR achieved by generated protein pockets with PocketGen aligns well with this practice, supporting its utility for stable and effective protein design.

In Table 1, the top-1, 3, 5, and 10 protein pockets generated by PocketGen (ranked by Vina score) consistently show the lowest Vina scores, achieving an average reduction of 0.476 compared to RFDiffusionAA. In addition to Vina scores, two other affinity metrics—MM-GBSA and GlideSP scores—further validate PocketGen’s ability to generate higher-affinity pockets, with reductions of 4.287 in MM-GBSA and 0.376 in GlideSP scores, respectively. Furthermore, PocketGen demonstrates competitive performance in pLDDT, scRMSD, and ΔscTM scores, underscoring its capability to produce high-affinity pockets while maintaining structural validity and sequence-structure consistency. With a 97% success rate in generating pockets with higher affinity than the reference cases (compared to a 93% success rate for the strongest baseline, RFDiffusionAA) on the CrossDocked dataset, PocketGen proves its effectiveness and applicability across diverse ligand molecules.

**Table 1.** The top 1/3/5/10 generated designable protein pocket (ranked by Vina score) on the CrossDocked dataset. The success rate measures the percentage of protein that the model can generate pockets with higher affinity than the reference ones in the datasets. Besides the Vina score, we additionally use MM-GBSA and min-in-space GlideSP score to evaluate the binding affinity. We report the average plDDT of the predicted pocket, the scRMSD of the pocket backbone coordinates, and the change of scTM scores of the whole protein. AF2 means the scores are calculated with AlphaFold2 as the folding tool (ESMFold results in Table S2). co indicates codesign, where codesign methods directly use the designed sequence for consistency calculation. The plDDT, scRMSD, and ΔscTM for PocketOpt are not reported, as PocketOpt keeps protein backbone structures fixed. We use 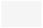 to mark the results of affinity-related metrics, 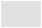 for pocket-structure related metrics, and 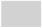 for whole protein structure metrics. We report the means and standard deviations over three independent runs with random seeds. The best results are indicated in **bold**.

|  | PocketOpt | DEPACT | dyMEAN | FAIR | RFDiffusion | RFAA | PocketGen |
| --- | --- | --- | --- | --- | --- | --- | --- |
| Top-1 generated protein pocket |  |  |  |  |  |  |  |
| Vina score (↓) | -9.216±0.154 | -8.527±0.061 | -8.540±0.107 | -8.792±0.122 | -9.037±0.080 | -9.216±0.091 | **-9.655±0.094** |
| MM-GBSA (↓) | -58.754±1.220 | -47.130±1.372 | 48.248±0.816 | -51.923±0.588 | -54.817±1.091 | -59.255±1.260 | **-63.542±0.717** |
| GlideSP (↓) | -8.612±0.127 | -7.495±0.053 | -7.472±0.088 | -7.584±0.094 | -8.485±0.069 | -8.540±0.065 | **-8.916±0.047** |
| Success Rate (↑) | 0.923±0.034 | 0.750±0.016 | 0.762±0.029 | 0.796±0.035 | 0.891±0.020 | 0.930±0.027 | **0.974±0.012** |
| pLDDT (AF2) (↑) | - | 82.164±0.241 | 83.053±0.397 | 83.285±0.240 | 84.432±0.152 | 86.571±0.178 | **86.830±0.145** |
| scRMSD (AF2) (↓) | - | 0.714±0.025 | 0.708±0.022 | 0.693±0.018 | 0.675±0.015 | 0.654±0.012 | **0.645±0.009** |
| ΔscTM (AF2) (↑) | - | -0.008±0.003 | -0.005±0.002 | -0.011±0.005 | 0.022±0.006 | 0.020±0.003 | **0.028±0.002** |
| ΔscTM (AF2+co) (↑) | - | -0.012±0.003 | -0.025±0.004 | -0.032±0.007 | - | - | **0.008±0.002** |
| Top-3 generated protein pockets |  |  |  |  |  |  |  |
| Vina score (↓) | -8.878±0.112 | -8.131±0.064 | -8.196±0.090 | -8.321±0.045 | -8.876±0.107 | -8.980±0.057 | **-9.353±0.063** |
| MM-GBSA (↓) | -53.372±1.164 | -43.790±1.029 | -44.151±0.534 | -46.050±0.809 | -52.423±0.847 | -53.593±0.722 | **-60.770±0.589** |
| GlideSP (↓) | -8.360±0.094 | -7.377±0.039 | -7.325±0.078 | -7.348±0.052 | -8.219±0.049 | -8.233±0.060 | **-8.670±0.056** |
| pLDDT (AF2) (↑) | - | 82.049±0.456 | 82.918±0.237 | 83.025±0.334 | 84.260±0.210 | **86.289±0.214** | 86.280±0.135 |
| scRMSD (AF2) (↓) | - | 0.713±0.017 | 0.722±0.011 | 0.692±0.016 | 0.685±0.007 | **0.659±0.014** | 0.660±0.012 |
| ΔscTM (AF2) (↑) | - | -0.011±0.004 | -0.006±0.002 | -0.008±0.003 | 0.021±0.003 | 0.022±0.002 | **0.026±0.003** |
| ΔscTM (AF2+co) (↑) | - | -0.016±0.005 | -0.026±0.004 | -0.034±0.003 | - | - | **0.005±0.001** |
| Top-5 generated protein pockets |  |  |  |  |  |  |  |
| Vina score (↓) | -8.702±0.090 | -7.786±0.052 | -7.974±0.049 | -7.943±0.035 | -8.510±0.073 | -8.689±0.044 | **-9.239±0.076** |
| MM-GBSA (↓) | -52.080±1.071 | -35.250±0.823 | -37.924±0.340 | -37.816±0.402 | -46.847±0.700 | -51.651±0.809 | **-58.083±0.561** |
| GlideSP (↓) | -8.173±0.089 | -7.126±0.035 | -7.294±0.042 | -7.289±0.041 | -8.022±0.030 | -8.093±0.048 | **-8.417±0.040** |
| pLDDT (AF2) (↑) | - | 82.445±0.307 | 82.763±0.102 | 83.748±0.271 | 84.505±0.288 | 85.617±0.105 | **85.969±0.080** |
| scRMSD (AF2) (↓) | - | 0.716±0.014 | 0.726±0.011 | 0.698±0.015 | 0.680±0.009 | 0.657±0.006 | **0.655±0.004** |
| ΔscTM (AF2) (↑) | - | -0.009±0.003 | -0.007±0.002 | -0.012±0.004 | 0.019±0.003 | 0.020±0.001 | **0.025±0.001** |
| ΔscTM (AF2+co) (↑) | - | -0.017±0.002 | -0.025±0.006 | -0.035±0.005 | - | - | **0.006±0.002** |
| Top-10 generated protein pockets |  |  |  |  |  |  |  |
| Vina score (↓) | -8.556±0.104 | -7.681±0.040 | -7.690±0.054 | -7.785±0.028 | -8.352±0.061 | -8.524±0.038 | **-9.065±0.057** |
| MM-GBSA (↓) | -49.257±0.821 | -32.534±0.680 | -33.118±0.269 | -33.670±0.440 | -45.726±0.830 | -47.325±0.540 | **-54.800±0.406** |
| GlideSP (↓) | -7.935±0.082 | -6.954±0.042 | -7.022±0.034 | -7.131±0.025 | -7.806±0.022 | -7.840±0.026 | **-8.196±0.027** |
| pLDDT (AF2) (↑) | - | 81.520±0.317 | 82.467±0.255 | 83.271±0.228 | 84.080±0.190 | 85.442±0.145 | **85.945±0.139** |
| scRMSD (AF2) (↓) | - | 0.712±0.013 | 0.733±0.014 | 0.706±0.013 | 0.688±0.009 | 0.680±0.010 | **0.659±0.007** |
| ΔscTM (AF2) (↑) | - | -0.014±0.002 | -0.006±0.001 | -0.010±0.003 | 0.016±0.002 | 0.019±0.001 | **0.023±0.002** |
| ΔscTM (AF2+co) (↑) | - | -0.018±0.004 | -0.030±0.002 | -0.033±0.002 | - | - | **0.004±0.002** |

To assess substructure validity and consistency with reference datasets, we conduct a qualitative substructure analysis (Table S4 and Figure S2). This analysis focuses on three covalent bonds in the residue backbone (C-N, C=O, and C-C), three dihedral angles in the backbone (*ϕ, ψ, ω* [58]), and four dihedral angles in the side chains (*χ*_1_, *χ*_2_, *χ*_3_, *χ*_4_ [59]). Following prior research [60, 61], we collect bond length and angle distributions from both the generated pockets and the test dataset and compute the Kullback-Leibler (KL) divergence to quantify the distance between these distributions. Lower KL divergence scores for PocketGen indicate its effectiveness in accurately replicating the geometric features observed in the reference data..

### Probing generative capabilities of PocketGen

Next, we explore PocketGen’s generative capabilities. Beyond designing high-quality protein pockets, generative models need to be efficient and maximize the yield of biochemical experiments—rapidly producing high-fidelity pocket candidates with only a small number of designs necessary to find a hit. Figure 3a compares the average **generation time** across various methods. Physics-based modeling (PocketOpt) and template-matching (DEPACT) can take over 1,000 seconds to generate 100 pockets. Advanced protein backbone generation models RFDiffusion and RFDiffusionAA are computationally expensive due to their diffusion-based architectures, requiring 1633.5 and 2210.1 seconds to design 100 pockets. Iterative refinement methods like PocketGen can significantly reduce generation time, with PocketGen taking just 44.2 seconds to generate 100 pockets.

**Figure 3.**
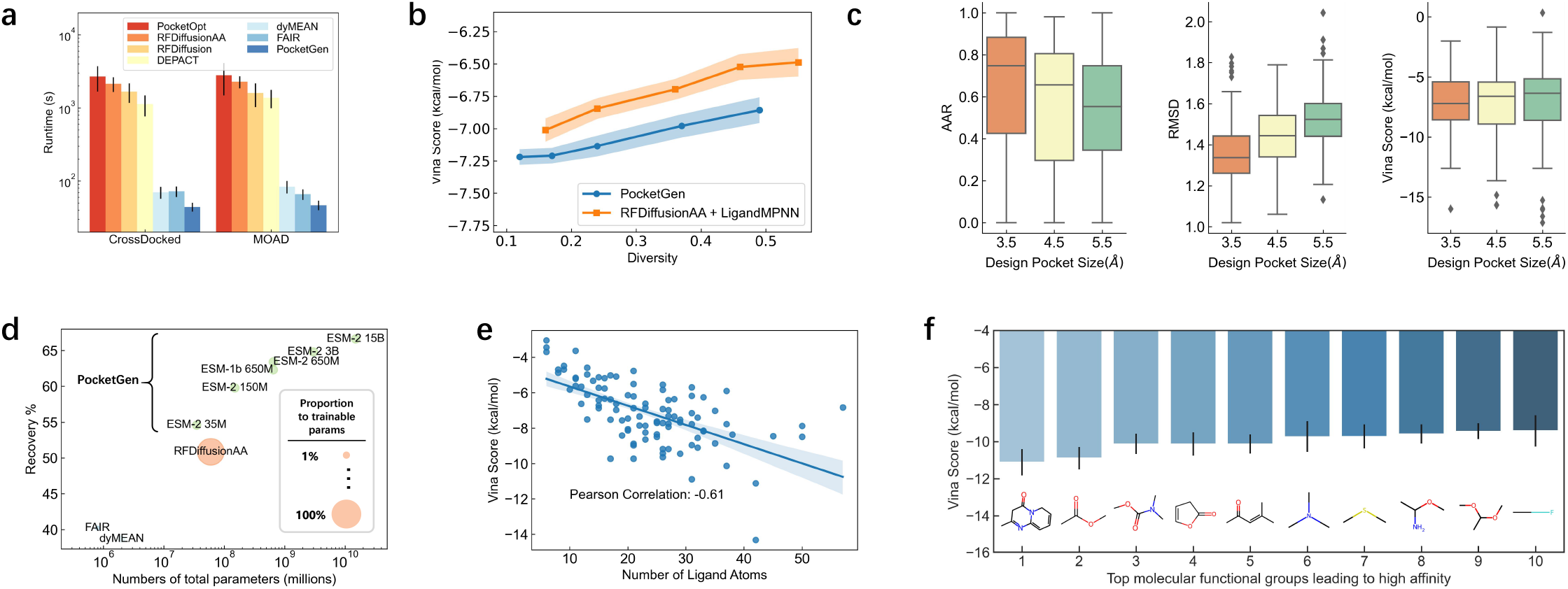
Exploring the capabilities of PocketGen. **a**, The average runtime of different methods for generating 100 protein pockets for a ligand molecule on the two benchmarks. Data are presented as mean values +/-standard deviation. The sample size for each method is 100. **b**, The trade-off between quality (measured by Vina score) and diversity (1-average pairwise sequence similarity) of PocketGen. We can balance the trade-off by tuning the temperate hyperparameter *τ*. We show the mean values with the standard deviations marked as shadows. **c**, The influence of the design pocket size on the metrics. We draw box plots and the sample size is 100. In box plots, the minimum is the smallest value, excluding outliers, marked at the end of the lower whisker. The first quartile (Q1), or 25th percentile, forms the lower edge of the box, while the median (50th percentile) is represented by a line within the box. The third quartile (Q3), or 75th percentile, forms the upper edge of the box. The maximum is the largest value, excluding outliers, marked at the end of the upper whisker. The whiskers extend to data points within 1.5 times the interquartile range (IQR), and any values beyond the whiskers are considered outliers. **d**, Performance w.r.t. model scales of pLMs using ESM series on CrossDocked dataset. The green dots represent PocketGen models with different ESMs. The bubble size is proportional to the number of trainable parameters. **e**, PocketGen tends to generate pockets with higher affinity for larger ligand molecules (Pearson Correlation *ρ* = −0.61, bands indicate 95% confidence interval). **f**, The top molecular functional groups leading to high affinity. The sample size is 100 and data are presented as mean values +/-standard deviation.

While recent methods for pocket generation focus on maximizing binding affinity with target molecules, this strategy may not always align with practical needs, where pocket diversity is equally important. Examining a batch of designed pockets, rather than a single design, improves the success rate of pocket design. Therefore, we investigate the relationship between binding affinity and the diversity of generated protein pockets in Figure 3b. **Diversity** is quantified as (−1 average pairwise pocket residue sequence similarity) and can be adjusted by altering the sampling temperature *τ* (with higher *τ* resulting in greater diversity). Figure 3b compares PocketGen with the most competitive baseline, RFDiffusionAA [16] + LigandMPNN [30], the latest version of ProteinMPNN [29]. We observe that there is a trade-off between binding affinity and diversity. PocketGen can generate protein pockets with higher affinity than RFDiffusionAA at the same level of diversity.

Figure 3c explores the effect of redesigned pocket size on PocketGen’s performance. The redesign process targets all residues with atoms within 3.5 Å, 4.5 Å, and 5.5 Å of any binding ligand atoms. We observe a slight decline in average AAR, RMSD, and Vina scores as the size of the redesigned pocket increases. This trend is likely due to the increased complexity and reduced contextual information in the case of larger redesigned pocket areas. Larger pockets tend to enable the exploration of structures with potentially higher affinity, as indicated by the lowest Vina scores, which reach -17.5 kcal/mol for designs with a 5.5 Å radius. This can be attributed to the enhanced structural complementarity in larger pocket designs. Extended Data Figure 1ab shows that PocketGen can generate full protein binders for two ligand molecules, with the generated protein binders achieving high scTM scores of 0.900 and 0.976.

A key feature that sets PocketGen apart from other pocket generation models is its integration of protein language models (pLMs). In addition to using ESM-2 650M [49] throughout our experiments, we evaluated a broader family of ESM models, ranging in model size from 8M to 15B trainable parameters. As shown in Figure 3d, PocketGen’s performance improves with the scaling of pLMs. Specifically, performance increases from 54.58% to 66.61% when transitioning from ESM-2 35M to ESM-2 15B models. This follows a logarithmic scaling law, consistent with trends observed in large language models [62]. PocketGen efficiently trains large pLMs by fine-tuning only the adapter layers while keeping most pLM layers fixed. As a result, PocketGen requires significantly fewer trainable parameters than RFDiffusionAA [16] (7.9M versus 82.9M trainable parameters).

The characteristics of the ligand molecule can affect the performance of PocketGen in generating binding pockets. Figure 3e shows the relationship between the average Vina score of generated pockets and the number of ligand atoms, revealing that PocketGen tends to create pockets with higher affinity for larger ligand molecules. This trend may result from the increased surface area for interaction, the presence of additional functional groups, and greater flexibility in the conformations of larger molecules [63, 64]. Key functional groups in ligand molecules that contribute to high binding affinity were identified using IFG [65]. Figure 3f highlights the top 10 molecular functional groups, which include hydrogen bond donors and acceptors (carbonyl groups), aromatic rings, sulfhydryl groups, and halogens. These groups facilitate favorable interactions with protein pockets, thereby enhancing binding affinity.

Since PocketGen also updates ligand structures during pocket generation, we use PoseBusters [66] to evaluate the structural validity of the updated ligands. A detailed validity check in Extended Data Figure 1e shows that PocketGen achieves over 95% across all tests in PoseBusters. This is expected, as PocketGen makes only minor updates to ligand structures during pocket generation, successfully maintaining ligand structural integrity. In Extended Data Figure 1c, we explore the relationship between binding affinity and the RMSD to the crystal structure in PDBBind. Using GIGN [100] to predict affinity (log K), we observe that generally, lower RMSD corresponds to higher affinity. Extended Data Figure 1d demonstrates that PocketGen improves most protein-ligand complexes in PDBBind by redesigning the binding pockets..

We conducted ablation studies (Table S5) and hyperparameter analysis (Figure S3) to assess the contribution of each module in PocketGen and the impact of hyperparameter choices on model performance. For comparison, we replaced the bilevel graph transformer in PocketGen with other popular encoders in structural biology, such as EGNN [67], GVP [68], and GMN [69]. The results indicate that the bilevel graph transformer and the integration of pLM into PocketGen significantly enhance performance. Furthermore, PocketGen demonstrates robustness to hyperparameter variations, consistently yielding competitive results.

### Generating protein pockets for therapeutic small molecules

We demonstrate PocketGen’s ability to redesign the pockets of antibodies, enzymes, and biosensors for specific target ligands, building upon previous research [3, 10, 16]. Specifically, we consider the following molecules: **Cortisol (HCY)** [70] is a primary stress hormone that raises glucose levels in the bloodstream and serves as a biomarker for stress and other conditions. We redesign the pocket of a cortisol-specific antibody (PDB ID 8cby), potentially aiding the development of immunoassays. **Apixaban (APX)** [71] is an oral anticoagulant approved by the FDA in 2012 for patients with non-valvular atrial fibrillation to reduce the risk of stroke and blood clots [72]. Apixaban targets Factor Xa (fXa) (PDB ID 2p16), an enzyme in blood coagulation that converts prothrombin into thrombin to facilitate clot formation. Redesigning the pocket of fXa has therapeutic implications. **Fentanyl (7V7)** [73] is a widely abused opioid contributing to the opioid crisis. Computationally designing fentanyl-binding proteins (biosensors) can support detection and neutralization efforts [10]. In Figure 4, PLIP [74] illustrates the interactions between the redesigned protein pockets and ligands, comparing these predicted interactions to the original binding patterns.

**Figure 4.**
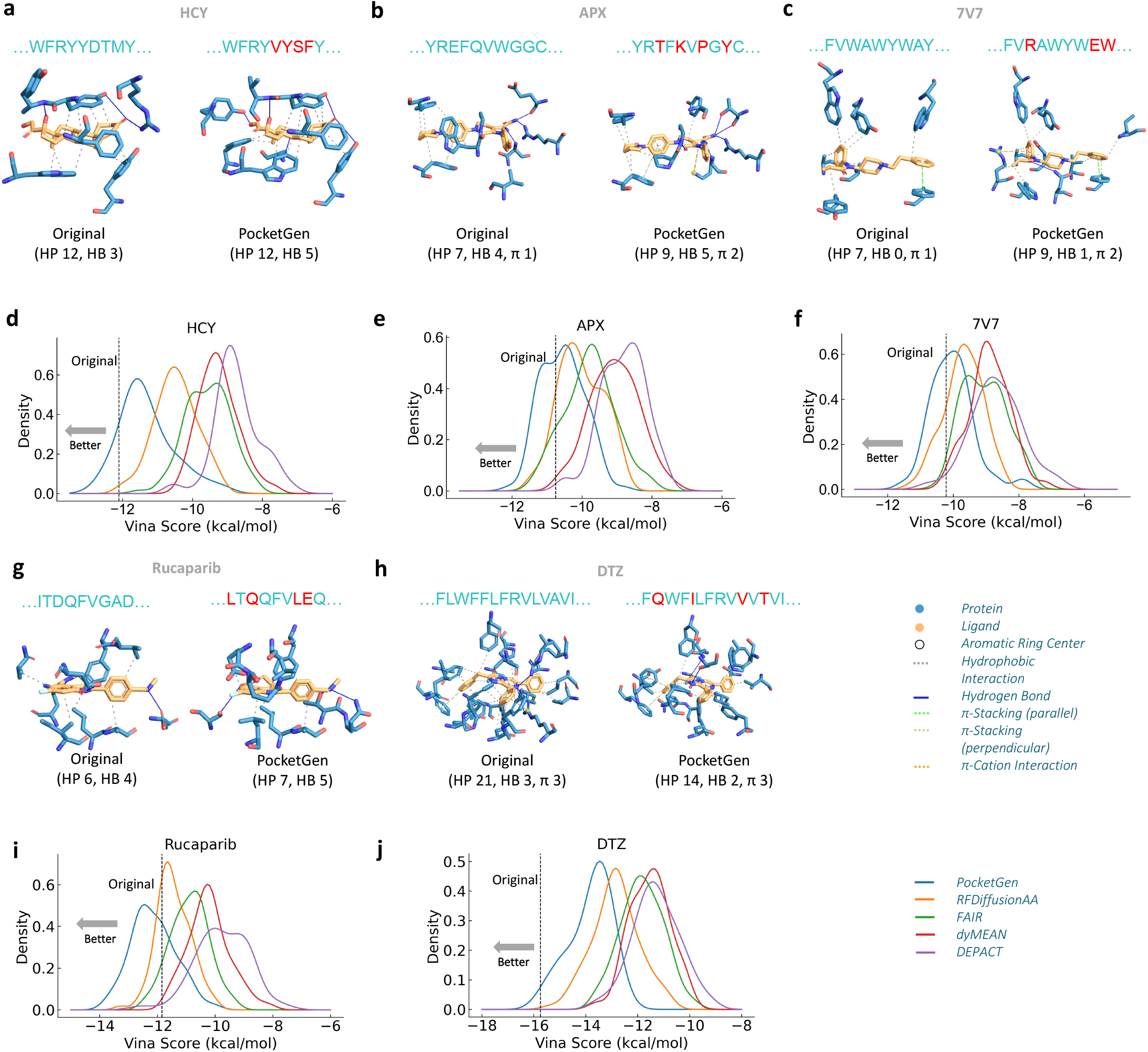
Using PocketGen to design protein pockets for binding with important ligands. **a, b, c**, Illustrations of protein-ligand interaction analysis for three target molecules (HCY, APX, and 7V7, respectively). ‘PocketGen’ refers to the protein pocket designed by PocketGen, and ‘Original’ denotes the original protein-ligand structure. ‘HP’ indicates hydrophobic interactions, ‘HB’ signifies hydrogen bonds, and ‘*π*’ denotes the *π*-stacking/cation interactions. In the residue sequences, red ones denote the designed residues that differ from the original pocket. **d, e, f**, The pocket binding affinity distributions of PocketGen and baseline methods for three target molecules (HCY, APX, and 7V7, respectively). We mark the Vina Score of the original pocket with the vertical dotted lines. For each method, we sample 100 pockets for each target ligand. The ratio of generated pockets by PocketGen with higher affinity than the corresponding reference pocket are 11%, 40%, and 45%, respectively. **g, h**, Protein-ligand interaction analysis for unseen proteins in the training dataset (PiB[21] and luxsit[8]). The target molecules are Rucaparib and DTZ, respectively. **i, j**, The pocket binding affinity distributions of PocketGen and baselines for Rucaparib and DTZ.

To generate pockets for the aforementioned small molecules, we pretrained PocketGen on the Binding MOAD dataset, excluding protein-ligand complexes considered in this analysis. The pockets produced by PocketGen successfully replicate most non-bonded interactions observed in experimentally measured protein-ligand complexes (achieving a 13/15 match for HCY) and introduce additional physically plausible interaction patterns not present in the original complexes. For example, the generated pockets for HCY, APX, and 7V7 molecules form 2, 3, and 4 extra interactions, respectively. Specifically for HCY, PocketGen preserves key interaction patterns such as hydrophobic interactions (TRP47, PHE50, TYR59, and TYR104) and hydrogen bonds (TYR59), while introducing two new hydrogen bond-mediated interactions within the pocket. For protein pockets designed to bind APX and 7V7 ligands, PocketGen maintains important interactions like hydrophobic contacts, hydrogen bonds, and *π*-*π* stacking while also establishing additional interactions—for example, a *π*-cation interaction with LYS192 for APX and hydrogen bonds with ASN35 for 7V7—thereby enhancing the binding affinity with the target ligands. PocketGen effectively captures non-covalent interactions derived from protein-ligand structure data while introducing new, plausible interaction patterns to optimize binding affinity.

With its ability to establish favorable protein-ligand interactions, PocketGen generates high-affinity pockets for these drug ligands. In Figure 4d,e,f, we present the affinity distributions of pockets generated by PocketGen compared to alternative methods. The ratio of generated pockets with higher affinity than the reference pocket is 11%, 40%, and 45% for PocketGen, respectively. In contrast, the best runner-up method, RFDiffusionAA, achieves only 0%, 10%, and 18% across the same cases. Protein stability is a critical factor in protein design, ensuring that the designed protein can fold into and maintain its three-dimensional structure [75]. Stability is quantified by the difference in Gibbs free energy (ΔΔ*G*) between the redesigned protein and the wild-type (original) protein, where ΔΔ*G* = Δ*G*_orig_ − Δ*G*_redesign_. A positive ΔΔ*G* value indicates increased stability, while a negative value suggests decreased stability. We used DDMut [76] to predict the change in stability for the pockets generated in Figure 4, with ΔΔ*G* values of 0.09 (HCY), 0.92 (APX), 0.13 (7V7), 0.27 (Rucaparib), and 0.02 (DTZ), respectively. These results suggest that PocketGen can generate protein structures likely to remain sufficiently stable to bind ligand molecules.

To demonstrate the generalization capability of PocketGen, we tested it on unseen proteins from the training set, including PiB [21] and luxsit [8], with the binding ligands Rucaparib and DTZ, respectively. Figures 4g and 4h show the interaction analysis, while Figures 4i and 4j present the distribution of Vina scores. PocketGen consistently outperforms other methods in generating higher-affinity pockets. Generating pockets with higher affinity for DTZ proved more challenging, as the original pocket was designed using site-saturation mutagenesis [8] to achieve optimal design. In Extended Data Figure 1f, we present case studies involving a pair of activity cliff ligand molecules (C19 and C52) [77] to further explore PocketGen’s adaptability. The generated interactions vary across molecular fragments: for one fragment, hydrogen bonds and hydrophobic interactions are generated, while for another fragment, halogen bonds are produced. This suggests that PocketGen has learned key protein-ligand interaction rules, allowing it to design high-affinity binding pockets.

### Interpreting protein-ligand interactions generated by PocketGen

We analyze attention maps learned by PocketGen using the generated pocket for the APX ligand. Figure 5a presents a 2D interaction plot drawn with the SchrÖdinger Maestro tool. To evaluate PocketGen’s recognition of key protein-ligand interactions, we plot the heatmap of attention weights produced by the final layer of its neural architecture. In Figure 5b, two attention heads are shown, with each row and column representing a protein residue or a ligand atom, respectively. The attention heatmaps are sparse, reflecting PocketGen’s use of sparse attention (Methods). The attention heads exhibit diverse patterns, focusing on different aspects of the interactions. For example, the first attention head emphasizes hydrogen bonds, assigning high weights to interactions between residue THR146, ASP220, and ligand atom 7. The second attention head captures *π*-*π* stacking and *π*-cation interactions, specifically between residue TYR99 and ligand atoms 15, 21, 23, 25, 29, and 33; and residue LYS192 and ligand atoms 1, 14, 17, 19, and 20. These findings suggest that, despite being data-driven, PocketGen has acquired biochemical knowledge to recognize intermolecular interactions.

**Figure 5.**
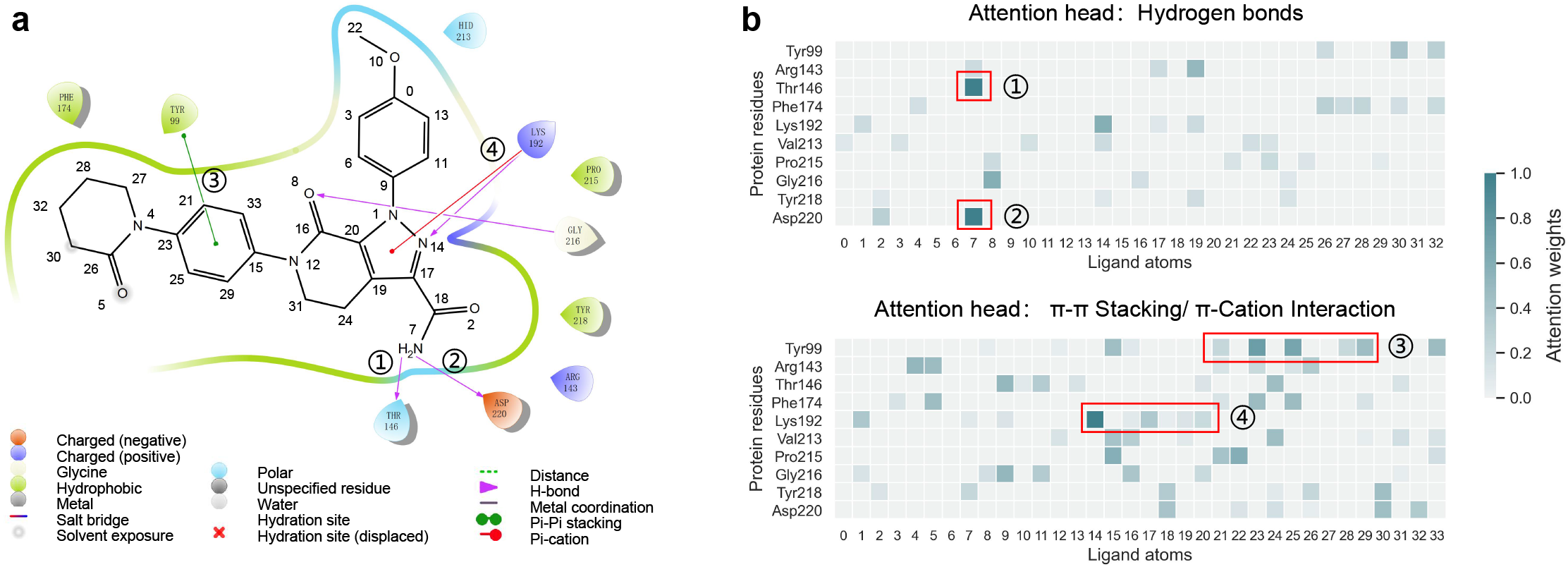
Attention maps in PocketGen capture interactions between atoms in protein and ligand molecules. **a**, The 2D interaction plot of the designed pocket by PocketGen for APX. **b**, The heatmap of attention matrices between residues and ligand atoms from the last layer of PocketGen. We show two selected attention heads with notable attention patterns marked with red rectangles. We notice that each head emphasizes different interactions. For example, PocketGen recognizes the **hydrogen bond** interaction and assigns a strong attention weight between residue ➀ THR146, ➁ ASP220, and ligand atom 7 in the first head. The *π***-***π* **stacking** and *π***-Cation** interactions of ➂ TYR99 and ➃ LYS192 are well captured in the second head. The values are normalized by the maximum value (*v*_max_) and the minimum value (*v*_min_) in each heatmap (i.e.,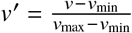).

## Discussion

Understanding how proteins bind to ligand molecules is critical for enzyme catalysis, immune recognition, cellular signal transduction, gene expression control, and other biological processes. Recent developments include deep generative models designed to study protein-ligand binding, like Lingo3DMol [78], ResGen [79], and PocketFlow [80] which generate *de novo* drug-like ligand molecules for fixed protein targets; NeuralPLexer [4] can create the structure of protein-ligand complexes given the protein sequence and ligand molecular graph. However, these models do not facilitate the *de novo* generation of protein pockets, the interfaces that bind with the ligand molecule for targeted ligand binding, critical in enzyme and biosensor engineering.

We developed PocketGen, a deep generative method capable of generating both the residue sequence and the full atom structures of the protein pocket region for binding with the target ligand molecule. PocketGen includes two main modules: a bilevel graph transformer for structural encoding and updates and a sequence refinement module that uses protein language models (pLMs) for sequence prediction. For structure prediction, the bilevel graph transformer directly updates the all-atom coordinates instead of separately predicting the backbone frame orientation and side-chain torsion angles. To achieve sequence-structure consistency and effectively leverage evolutionary knowledge from pLMs, a structural adapter is integrated into protein language models for sequence updates. This adapter employs cross-attention between sequence and structure features to promote information flow and ensure sequence-structure consistency. Extensive experiments across benchmarks and case studies involving therapeutic ligand molecules illustrate PocketGen’s ability to generate high-fidelity pocket structures with high binding affinity and favorable interactions with target ligands. Analysis of PocketGen’s performance across various settings reveals its proficiency in balancing diversity and affinity and generalizing across different pocket sizes. Additionally, PocketGen offers computational efficiency, significantly reducing runtime compared to traditional physics-based methods, making it feasible to sample large quantities of pocket candidates. PocketGen surpasses existing methods in efficiently generating high-affinity protein pockets for target ligand molecules, finding important interactions between atoms on protein and ligand molecules, and attaining consistency in sequence and structure domains.

PocketGen creates several fruitful directions for future work. PocketGen could be expanded to design larger areas of the protein beyond the pocket area. While PocketGen has been evaluated on larger pocket designs, modifications will be required to enhance scalability and robustness for generating larger protein areas. Another fruitful future direction involves incorporating additional biochemical priors, such as subpockets [81] and interaction templates [17], to improve generalizability and success rates. For instance, despite overall dissimilarity, two protein pockets might still bind the same fragment if they share similar subpockets [82]. Moreover, conducting wet lab experiments could provide empirical validation of PocketGen’s effectiveness. Approaches such as PocketGen have the potential to advance areas of machine learning and bioengineering and help with the design of small molecule binders and enzymes.

## Methods

### Overview of PocketGen

Unlike previous methods focusing on protein sequence or structure generation, we aim to co-design both residue types (sequences) and 3D structures of the protein pocket that can fit and bind with target ligand molecules. Inspired by previous works on structure-based drug design [79, 81] and protein generation [34, 35], we formulate pocket generation in PocketGen as a conditional generation problem that generates the sequences and structures of pocket conditioned on the protein scaffold (other parts of the protein except the pocket region) and the binding ligand. To be specific, let 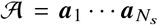 denote the whole protein sequence of residues, where *N*_*s*_ is the length of the sequence. The 3D structure of the protein can be described as a point cloud of protein atoms 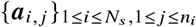 and let ***x*** (***a***_*i, j*_) ∈ ℝ^3^ denote the 3D coordinate of protein atoms. *n*_*i*_ is the number of atoms in a residue determined by the residue types. The first four atoms in any residue correspond to its backbone atoms (*C*_*α*_, *N, C, O*), and the rest are the side-chain atoms. The ligand molecule can also be represented as a 3D point cloud 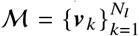 where ***v***_*k*_ denotes the atom feature. Let ***x*** (***v***_*k*_) denotes the 3D coordinates of atom ***v***_*k*_. Our work defines the protein pocket as a set of residues in the protein closest to the binding ligand molecule: ℬ = ***b***_1_ · · · ***b***_*m*_. The pocket ℬ can thus be represented as an amino acid subsequence of a protein: 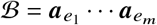 where ***e*** = {*e*_1_, · · ·, *e*_*m*_} is the index of the pocket residues in the whole protein. The index ***e*** can be formally given as: 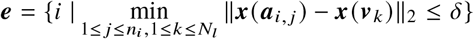, where ∥·∥_2_ is the *L*_2_ distance norm and *δ* is the distance threshold. According to the distance range of pocket-ligand interactions [45], we set *δ* = 3.5 A in the default setting. With the above-defined notations, PocketGen aims to learn a conditional generative model formally defined as :

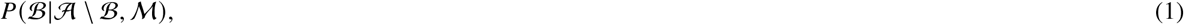

where 𝒜 \ ℬ denotes the other parts of the protein except the pocket region. We also adjust the structure ligand molecule ℳ in PocketGen to encourage protein-ligand interactions and reduce steric clashes.

To effectively generate the structure and the sequence of the protein pocket ℬ, the equivariant bilevel graph transformer and the sequence refinement module with pretrained protein language models and adapters are proposed, which will be discussed in the following paragraphs. The illustrative workflow is depicted in Fig. 1.

### Equivariant bilevel graph transformer

It is critical to model the complex interactions in the protein pocket-ligand complexes for pocket generation. However, the multi-granularity (e.g., atom-level and residue-level) and multi-aspect (intra-protein and protein-ligand) nature of interactions brings a lot of challenges. Inspired by recent works on hierarchical graph transformer [81] and generalist equivariant transformer [83], we propose a novel equivariant bilevel graph transformer to well model the multi-granularity and multi-aspect interactions. Each residue or ligand is represented as a block (i.e., a set of atoms) for the conciseness of representation and ease of computation. Then the protein-ligand complex can be abstracted as a geometric graph of sets 𝒢 = (𝒱,ℰ), where 𝒱= {***H***_*i*_, ***X***_*i*_ |1≤ *i* ≤ *B*} denotes the blocks and ℰ= {*e*_*i j*_ |1 ≤ *i, j* ≤ *B*} include all the edges between blocks (*B* is the total number of blocks). We added self-loops to the edges to capture interactions within the block (e.g., the interactions between ligand atoms). Our model adaptively assigns different numbers of channels to ***H***_*i*_ and ***X***_*i*_ to accommodate different numbers of atoms in residues and ligands. For example, given a block with *n*_*i*_ atoms, the corresponding block has ***H***_*i*_ 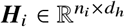 indicating the atom features (*d*_*h*_ is the feature dimension size) and 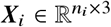 denoting the atom coordinates. Specifically, the *p*-th row of *H*_*i*_ and ***X***_*i*_ corresponds to the *p*-th atom’s trainable feature (i.e., ***H***_*i*_ [*p*]) and coordinates (i.e., ***X***_*i*_ [*p*]) respectively. The trainable feature ***H***_*i*_[*p*] is first initialized with the concatenation of atom type embedding, residue/ligand embeddings, and the atom positional embeddings. To build ℰ, we connect the *k*-nearest neighboring residues according to the pairwise *C*_*α*_ distances. To reflect the interactions between the protein pocket and ligand, we add edges between all the pocket residue and the ligand block. We describe the modules in PocketGen’s equivariant bilevel graph transformer, bilevel attention module, and equivariant feed-forward networks.

### Bilevel attention module

Our model captures both **atom-level** and **residue/ligand-level** interactions with the bilevel attention module. Firstly, given two block *i* and *j* connected by an edge *e*_*i j*_, we obtain the query, the key, and the value matrices with the following transformations:

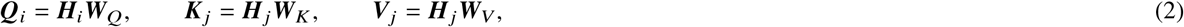

where 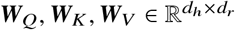 are trainable parameters.

To calculate the **atom-level attention** across the *i*-th and *j* -th block, we denote 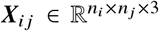 and 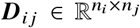 as the relative coordinates and distances between atom pairs in block *i* and *j*, namely, ***X***_*i j*_ [ *p, q*] = ***X***_*i*_ [ *p*] − ***X*** _*j*_ [*q*], ***D***_*i j*_ [ *p, q*] = ∥ ***X***_*i j*_ [ *p, q*] ∥_2_. Then we have:

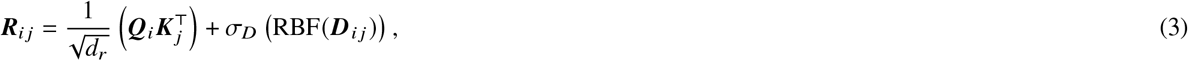

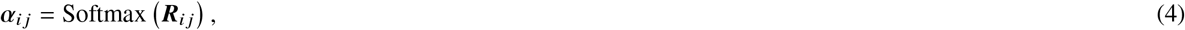

where *σ*_*D*_(·) is a Multi-Layer Perceptron (MLP) that adds distance bias to the attention calculation. RBF embeds the distance with radial basis functions. 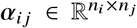 is the atom-level attention matrix obtained by applying row-wise Softmax on 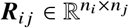. To encourage sparsity in the attention matrix, we keep the top-*k*^′^ elements of each row in ***α***_*i j*_ and set the others as zeros.

The **residue/ligand-level attention** from the *j* -th block to the *i*-th block is calculated as:

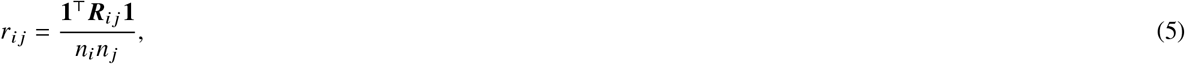

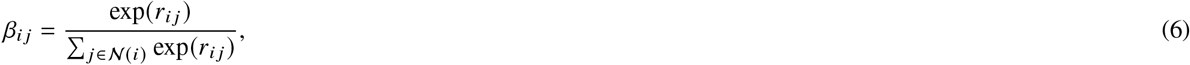

where **1** refers to the column vector with all elements set as ones and 𝒩 (*i*) denotes the neighboring blocks of *i. r*_*i j*_ sums up all values in ***R***_*i j*_ to represent the overall correlation between blocks *i* and *j*. Subsequently, *β*_*i j*_ denotes the attention across blocks at the block level.

We can update the representations and coordinates using the above atom-level and the residue/ligand-level attentions. PocketGen only updates the coordinates of the residues in the pocket and the ligand molecule. The other protein residues are fixed. Specifically, for the *p*-th atom in block *i*:

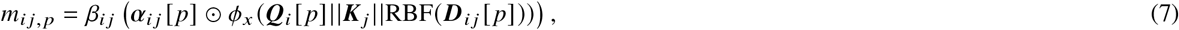

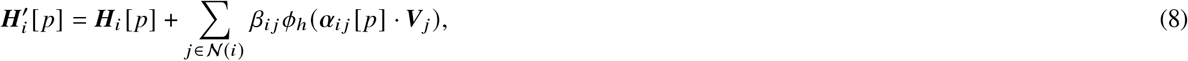

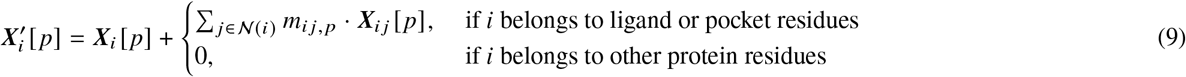

where *ϕ*_*h*_ and *ϕ*_*x*_ are MLPs with concatenated representations as input (concatenation along the second dimension and ***Q***_*i*_[*p*] is repeated along rows). ⊙ computes the element-wise multiplication. 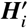 and 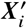 denote the updated representation and coordinate matrices, and we can verify that the dimension size of 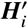 and 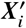 remains the same regardless of the neighboring block size *n*_*j*_. Furthermore, as the attention coefficients ***α***_*i j*_ and *β*_*i j*_ are invariant under E(3) transformations, the modification of 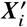 adheres to E(3)-equivariance. Additionally, the permutation of atoms within each block does not affect this update process.

### Equivariant feed-forward network

We adapted the feed-forward network module (FFN) in the transformer model[84] to update ***H***_*i*_ and ***X***_*i*_. Specifically, the representation and coordinates of atoms are updated to consider the block’s feature/geometric centroids (means). The centroids are denoted as:

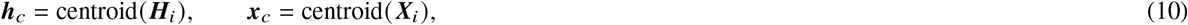

Then we obtain the relative coordinate Δ***x*** _*p*_ and the relative distance representation ***r*** _*p*_ based on the L2 norm of Δ***x*** _*p*_:

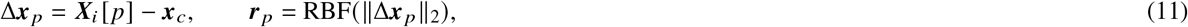

The representation and coordinates of atoms are updated with MLPs *σ*_*h*_ and *σ*_*x*_. The centroids are integrated to inform of the context of the block:

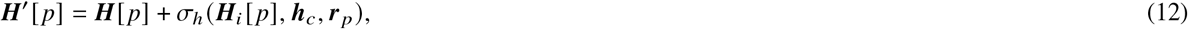

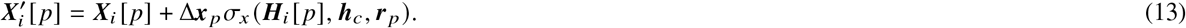

To stabilize and accelerate training, layer normalization [85] is appended at each layer of the equivariant bilevel graph transformer to normalize ***H***. The equivariant feed-forward network satisfies E(3)-equivariance. Thanks to each module’s E(3)-equivariance, the whole proposed bilevel graph transformer has the desirable property of E(3)-equivariance (Theorem 1 in Supplementary Information shows the details). In PocketGen, we use E(3) equivariant model for its simplicity similar to previous works[86, 87], which is capable enough to achieve strong performance. We are aware that an SE(3) equivariant model architecture would be better for learning the chirality-related properties of the protein, which we left for future exploration.

### Sequence refinement with protein language models and adapters

Protein language models (pLMs), such as the ESM family of models [40, 41], have learned extensive evolutionary knowledge from the vast array of natural protein sequences, demonstrating a strong ability to design protein sequences. In PocketGen, we propose to leverage pLMs to help refine the designed protein pocket sequences. To infuse the pLMs with structural information, we implant lightweight structural adapters inspired by previous works [88, 89]. Different from LM-Design [89] which focuses on protein sequence design given fixed backbone structure, PocketGen codesigns both the amino acid sequence as well as the full atom structure of the protein pocket. In our default setting, only one structural adapter was placed after the last layer of pLM. Only the adapter layers are fine-tuned during training, and the other layers of PLMs are frozen to save computation costs. The structural adapter mainly has the following two parts.

### Structure-sequence cross attention

The structural representation of the *i*-th residue 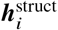 is obtained by mean pooling of ***H***_*i*_ from the bilevel graph transformer. In the input to the pLMs, the pocket residue types to be designed are assigned with the mask, and we denote the *i*-th residue representation from pLMs as 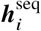. In the structural adapter, we perform cross-attention between the structural representations 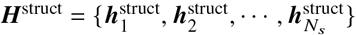 and sequence representations 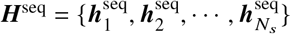.

The query, key, and value matrices are obtained as follows:

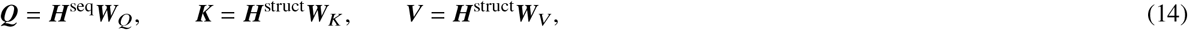

where 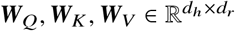 are trainable weight matrices. Rotary positional encoding [90] is applied to the representations, and we omit it in the equations for simplicity. The output of the cross attention is obtained as:

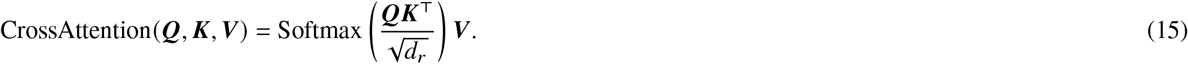

### Bottleneck feed-forward network

A bottleneck feed-forward network (FFN) is appended after the cross-attention to impose non-linearity and abstract representations, inspired by previous works such as Houlsby et al.[88]. The intermediate dimension of the bottleneck FFN is set to be half of the default representation dimension. Finally, the predicted pocket residue type ***p***_*i*_ is obtained using an MLP on the output residue representation.

### Training protocol

Inspired by AlphaFold2 [39], we use a recycling strategy for model training. Recycling facilitates the training of deeper networks without incurring extra memory costs by executing multiple forward passes and computing gradients solely for the final pass. The training loss of PocketGen is the weighted sum of the following three losses:

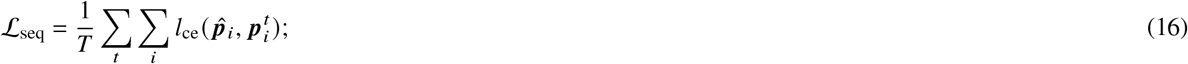

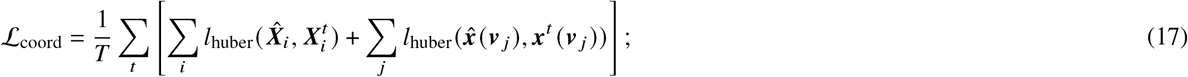

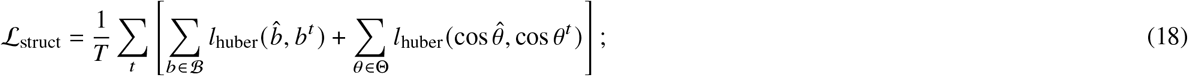

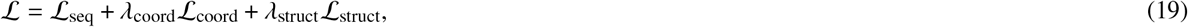

where *T* is the total refinement rounds.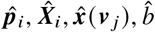, and cos 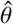 are the ground-truth residue types, residue coordinates, and ligand coordinates, bond lengths, and bond/dihedral angles; 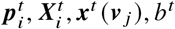, *b*^*t*^, and cos *θ*^*t*^ are the predicted ones at the *t*-th round by PocketGen. The sequence loss ℒ_seq_ is the cross-entropy loss for pocket residue type prediction; the coordinate loss ℒ_coord_ uses huber loss [91] for the training stability; the structure loss ℒ_struct_ is added to supervised bond lengths and bond/dihedral angles for realistic local geometry. ℬ and Θ denote all the bonds and angles in the protein pocket (including side chains). *λ*_coord_, and *λ*_struct_ are hyperparameters balancing the three losses. We perform a grid search over {0.5, 1.0, 2.0, 3.0} and choose these hyperparameters based on the validation performance to select the specific parameter values. In the default setting, we set *λ*_coord_ to 1.0 and *λ*_struct_ to 2.0.

### Generation protocol

In the generation procedure, PocketGen initializes the sequence with uniform distributions over 20 amino acid types and the coordinates based on linear interpolations and extrapolations. Specifically, we initialize the residue coordinates with linear interpolations and extrapolations based on the nearest residues with known structures in the protein. Denote the sequence of residues as 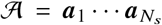, where *N*_*s*_ is the length of the sequence. Let ***x a***_*i*,1_ ∈ ℝ^3^ denote the *C*_*α*_ coordinate of the *i*-th residue. We take the following strategies to determine the *C*_*α*_ coordinate of the *i*-th residue: (1) We use linear interpolation if there are residues with known coordinates at both sides of the *i*-th residue. Specifically, assume *p* and *q* are the indexes of the nearest residues with known coordinates at each side of the *i*-th residue (*p* < *i* < *q*), we have: 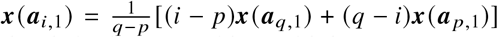. (2) We conduct linear extrapolation if the *i*-th residue is at the ends of the chain, i.e., no residues with known structures at one side of the *i*-th residue. Specifically, let *p* and *q* denote the index of the nearest and the second nearest residue with known coordinates. The position of the *i*-th residue can be initialized as 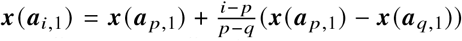. Inspired by previous works [33, 34], we initialize the other backbone atom coordinates according to their ideal local coordinates relative to the *C*_*α*_ coordinates. We initialize the side-chain atoms’ coordinates with the coordinate of their corresponding *C*_*α*_, added with Gaussian noise. We initialize the ligand molecular structure with the reference ligand structure from the dataset. The ligand structure is updated during pocket generation and the updated ligand is used for Vina score calculation.

Since the number of pocket residue types and the number of side chain atoms are unknown at the beginning of generation, each pocket residue is assigned 14 atoms, the maximum number of atoms for residues. After rounds of refinement by PocketGen, the pocket residue types are predicted, and the full atom coordinates are determined by mapping the coordinates to the predicted residue types (taking the first n coordinates according to residue type). In PocketGen, we directly predict the absolute atom coordinates, which reduces the model complexity and flexibly captures atom interactions. We also notice PocketGen aligns with the recent trend of directly predicting full atom coordinates. For example, the recent AlphaFold3 [92] directly predicts the full atom coordinates, replacing the AlphaFold2 structure module that operated on amino-acid-specific frames and side-chain torsion angles, and achieves better performance on protein structure prediction. For generation efficiency, we set the number of refinement rounds to 3.

### Experimental setting

#### Datasets

We consider two widely used datasets for benchmark evaluation: **CrossDocked** dataset [42] contains 22.5 million protein-molecule pairs generated through cross-docking. Following previous works [24, 60, 93], we filter out data points with binding pose RMSD greater than 1 Å, leading to a refined subset with around 180k data points. For data splitting, we use mmseqs2 [94] to cluster data at 30% sequence identity, and randomly draw 100k protein-ligand structure pairs for training and 100 pairs from the remaining clusters for testing and validation, respectively; **Binding MOAD** dataset [43] contains around 41k experimentally determined protein-ligand complexes. Following previous work [95], we keep pockets with valid and moderately ‘drug-like’ ligands with QED score ≥0.3. We further filter the dataset to discard molecules containing atom types ∉ {*C, N, O, S, B, Br, Cl, P, I, F*} as well as binding pockets with non-standard amino acids. Then, we randomly sample and split the filtered dataset based on the Enzyme Commission Number (EC Number) [44] to ensure different sets do not contain proteins from the same EC Number main class. Finally, we have 40k protein-ligand pairs for training, 100 pairs for validation, and 100 pairs for testing. For all the benchmark tasks in this paper, PocketGen and all the other baseline methods are trained with the same data split for a fair comparison. In real-world pocket generation and optimization case studies, the protein structures were downloaded from PDB [96].

#### Implementation

Our PocketGen model is trained with Adam [97] optimizer for 5k iterations, where the learning rate is 0.0001, and the batch size is 64. We report the results corresponding to the checkpoint with the best validation loss. It takes around 48 hours to finish training on 1 Tesla A100 GPU from scratch. In PocketGen, the number of attention heads is set as 4; the hidden dimension d is set as 128; *k* is set to 8 to connect the *k*-nearest neighboring residues to build ℰ; *k*^′^ is set as 3 to encourage sparsity in the attention matrix. For all the benchmark tasks of pocket generation and optimization, PocketGen and all the other baseline methods are trained with the same data split for a fair comparison. We follow the implementation codes provided by the authors to obtain the results of baseline methods. Algorithm 1 and 2 in the supplementary show the pseudo-codes of the training and generation process of PocketGen.

#### Baseline methods

PocketGen is compared with five state-of-the-art representative baseline methods. **PocketOptimizer** [18] is a physics-based method that optimizes energies such as packing and binding-related energies for ligand-binding protein design. Following the suggestion of the paper, we fixed the backbone structures. **DEPACT** [17] is a template-matching method that follows a two-step strategy [98] for pocket design. It first searches the protein-ligand complexes in the database with similar ligand fragments. It then grafts the associated residues into the protein scaffold to output the complete protein structure with PACMatch [17]. Both the backbone and the sidechain structures are changed in DEPACT. **RFDiffusion**[26], **RFDiffusionAA**[16], **FAIR**[24], and **dyMEAN**[25] are deep-learning-based models that for protein generation. RFDiffusion does not explicitly model protein-ligand interactions and is not directly applicable to small molecule-binding protein generation.

Following the suggestions in RFDiffusion[26] and RFDiffusionAA [16], we use a heuristic attractive-repulsive potential to encourage the formation of pockets with shape complementarity to a target molecule. The residue sequence for the generated protein by RFDiffusion is derived with ProteinMPNN, and the side-chain conformation is decided with Rosetta[99] side-chain packing. RFDiffusionAA is the latest version of RFDiffusion, which can directly generate protein structures surrounding small molecules by combining residue-based representation of amino acids with atomic representation of small molecules. For RFDiffusion and RFDiffusionAA, we let them in paint the pocket area to obtain a consistent setting with other methods for comparison. We also note that RFDiffusion and RFDiffusionAA do not provide the training/finetuning scripts, so we use the provided pre-trained checkpoints for all the related experiments in our paper. FAIR [24] was specially designed for full-atom protein pocket design via iterative refinement. dyMEAN[25] was originally proposed for full atom antibody design, and we adapted it to our pocket design task with proper modifications. Detailed information on baselines is included in Supplementary Notes. The setting of the key hyperparameters is summarized in Table. S6. All the baselines are run on the same Telsa A100 GPU for a fair comparison with our PocketGen.

## Data availability

This study’s training and test data are available at Zenodo [104]. The project website for PocketGen is at https://zitniklab.hms.harvard.edu/projects/PocketGen.

## Code availability

The source code of this study is freely available at GitHub (https://github.com/zaixizhang/PocketGen) and can be accessed via DOI [103].

## Acknowledgements

Z.X.Z. gratefully acknowledges the support of grants from the National Natural Science Foundation of China (No. 623B2095) and the Excellent PhD Students Overseas Study Program of the University of Science and Technology of China. Q.L. gratefully acknowledges the support of grants from the National Natural Science Foundation of China (No. 62337001) and the Fundamental Research Funds for the Central Universities. M.Z. gratefully acknowledges the support of NIH R01-HD108794, NSF CAREER 2339524, US DoD FA8702-15-D-0001, awards from Harvard Data Science Initiative, Amazon Faculty Research, Google Research Scholar Program, AstraZeneca Research, Roche Alliance with Distinguished Scientists, Sanofi iDEA-iTECH Award, Pfizer Research, Chan Zuckerberg Initiative, John and Virginia Kaneb Fellowship award at Harvard Medical School, Biswas Computational Biology Initiative in partnership with the Milken Institute, and Kempner Institute for the Study of Natural and Artificial Intelligence at Harvard University. We thank Dr. Enhong Chen, Dr. Yaoxi Chen, and Dr. Haiyan Liu from the University of Science and Technology of China for their constructive discussions on implementing and evaluating baseline methods, which greatly helped this research.

## Author contributions statement

Z.X.Z., Q.L., and M.Z. designed the research, Z.X.Z. conducted the experiments, Z.X.Z., Q.L., and M.Z. analyzed the results. Z.X.Z., W.X.S., Q.L., and M.Z. wrote the manuscript. All authors reviewed the manuscript.

## Figure Legends/Captions

**Extended Data Figure 1:**
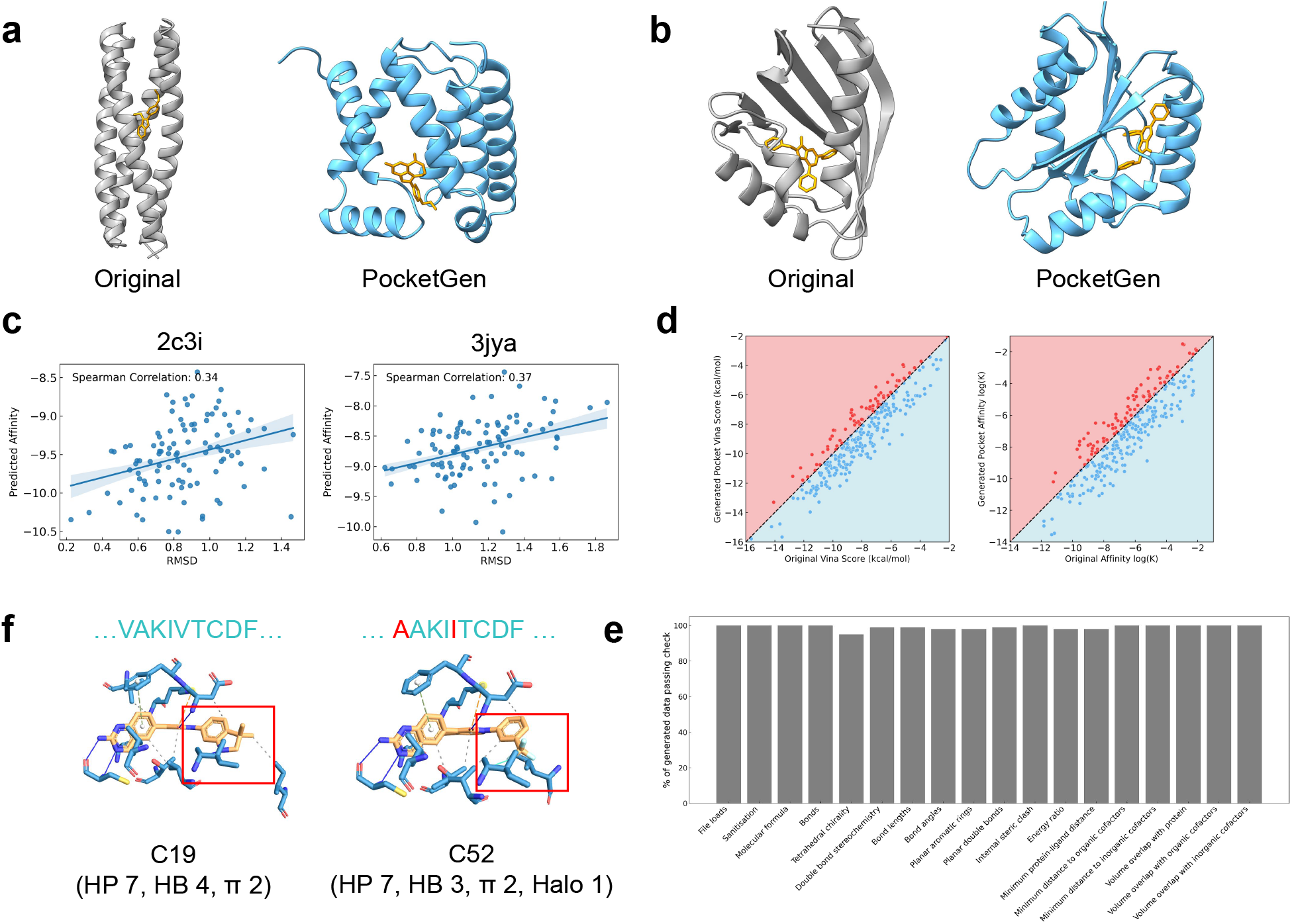
More case studies and evaluations of PocketGen. **a**, The originally designed protein binder for Rucaparib [21](left panel) and the generated protein binder by PocketGen (right panel). **b**, The originally designed protein binder for DTZ [8](left panel) and the generated protein binder by PocketGen (right panel). Note that PocketGen generates the whole protein instead of the pocket region in **a&b**. The generated protein binder has high scTM scores (0.900 and 0.976). **c**, The predicted affinity (log K) by GIGN [100] of the generated pockets by PocketGen with respect to RMSD. We randomly select two protein-ligand complexes from PDBBind (PDB id 2c3i and 3jya). **d**, The Vina score/binding affinity (log K) of the generated pockets by PocketGen and the original pockets from PDBBind. The black region/dots indicate the generated pockets have higher affinities than the original pockets while the red region/dots indicate lower affinities. **f**, The generated interactions by PocketGen with respect to a pair of activity cliff ligand molecules, i.e., C19 and C52 [77]. As marked with red rectangles, PocketGen adaptively generates different interactions for different molecular fragments (hydrogen bonds+hydrophobic interactions and halogen bonds respectively). ‘HP’ indicates hydrophobic interactions, ‘HB’ signifies hydrogen bonds, ‘*π*’ denotes the *π*-stacking/cation interactions, and ‘Halo’ indicates the Halogen bonds. **e**, Detailed validity check with PoseBusters on CrossDocked and Binding MOAD.

## Supplementary Information

### Supplementary Notes

#### Training and Generation Algorithms

##### Algorithm 1 Training Algorithm of PocketGen

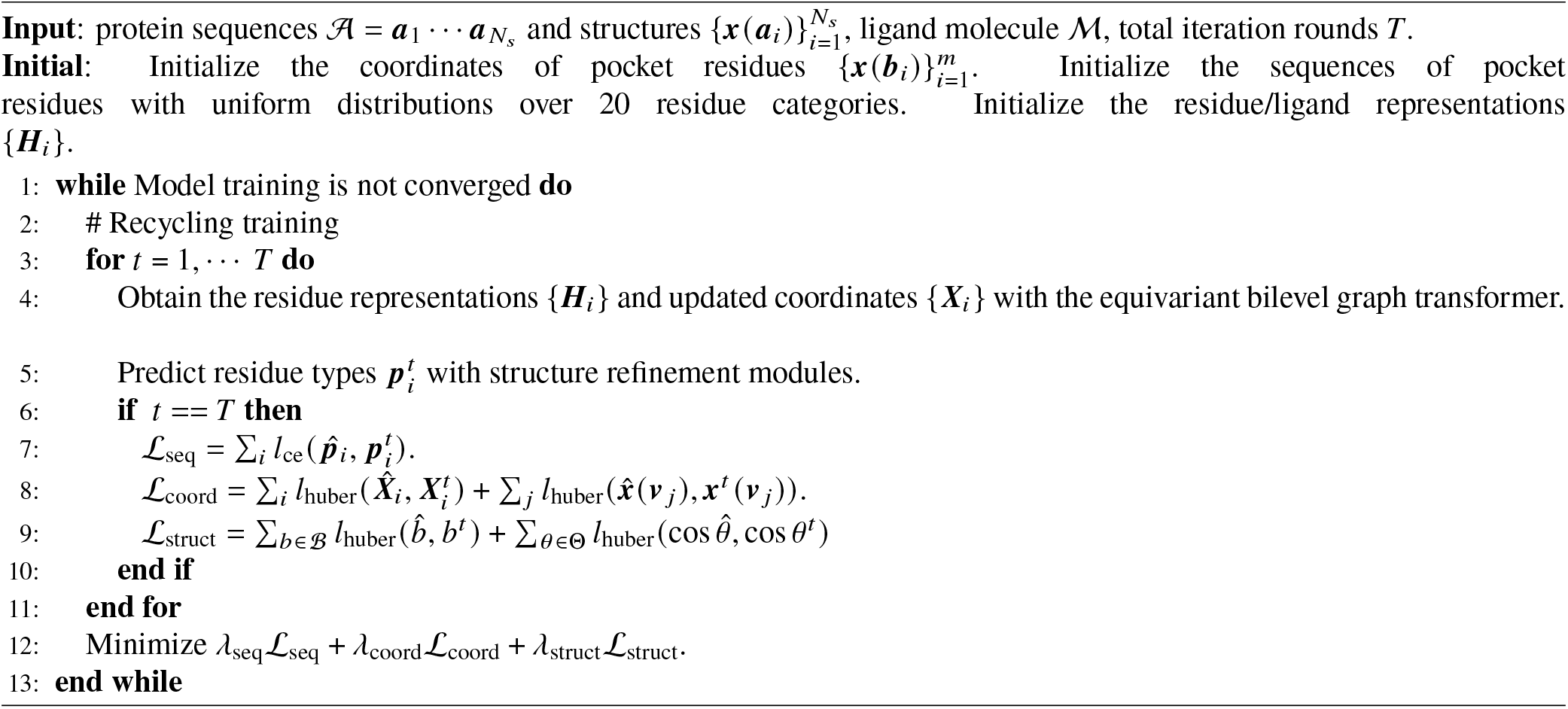

##### Algorithm 2 Generation Algorithm of PocketGen

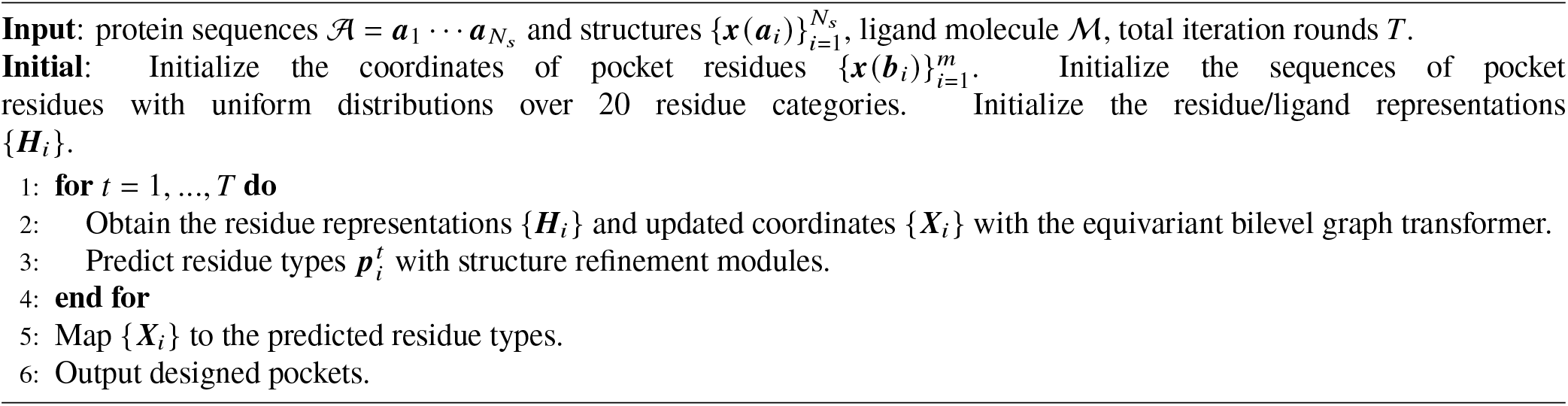

#### Loss Functions

In Equation 19, we use Huber loss [91] in ℒ_coord_ and ℒ_struct_ for the stability of optimization, which are defined as follows:

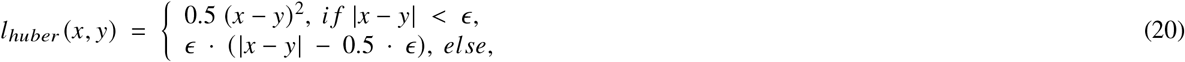

where *x* and *y* represent the predicted and ground-truth coordinates/ bond length/ angles. The Huber loss has the following property: if the L1 norm of |*x*−*y*| is smaller than *ϵ*, it is MSE loss, otherwise it is L1 loss. At the beginning of the model training, the deviation between the predicted and ground-truth coordinates is large, and the L1 term makes the loss less sensitive to outliers than MSE loss. The deviation is small when the training is almost complete, and the MSE loss is applied for further finetuning. In practice, we find that directly using MSE loss sometimes leads to instability at the beginning of the training (e.g., very large gradient norm), while Huber loss makes the training procedure more stable. Following the suggestion of previous works [32, 34], we set *ϵ* = 1 in all our experiments.

#### Implementation Details of Baseline Methods and Hyperparameter Settings

**PocketOptimizer** [18] ^1^ is a physic-based computational protein design method that predicts mutations in the binding pockets of proteins to increase affinity for a specific ligand. We use the latest version, i.e., PocketOptimizer 2.0 [18]. There are generally four main steps in PocketOptimizer: structure preparation, flexibility sampling, energy calculations, and computation of design solutions. Specifically for the energy calculations, both packing-related energies and binding-related energies are considered. Following the suggestions in the original paper, we use AMBER ff14S force field [101] for energy computation and the Dunbrack rotamer library [102] for rotamer sampling for PocketOptimizer in our implementation. We fixed the backbone structures following the suggestions of the original paper. As for the output design solutions, we select the top 100 designs identified by PocketOptimizer based on integer linear programming for downstream metric calculations.

**DEPACT** [17] ^2^ is a template-matching method that follows a two-step strategy for pocket design. Firstly, it searches the protein-ligand complexes in the template database with similar ligand fragments and constructs a cluster model (a set of pocket residues). The template databases are constructed based on the corresponding training datasets for fair comparisons. Secondly, it grafts the cluster model into the protein pocket with PACMatch. It works by placing residues from the cluster model on protein scaffolds by matching the atoms of residues with atoms of the protein scaffold. The backbone coordinates of the pocket residues are also modified in the process. The qualities of the generated pockets are evaluated and ranked based on a statistical scoring function. We take the top 100 designed pockets for evaluation. The output of DEPACT+PACMatch is complete protein structures with redesigned pockets. In the paper, we only use DEPACT to represent the whole method of DEPACT+PACMatch for conciseness.

**RFDiffusion** [26] ^3^ is one of the state-of-the-art method for *de novo* protein backbone generation. It combines the RoseTTAFold structure prediction network with the diffusion probabilistic models (DDPMs) framework. To model the influence of ligand molecules, we use a heuristic attractive-repulsive potential to encourage the formation of pockets with shape complementarity to a target molecule following the suggestions of RFDiffusion[26] and RFDiffusionAA [16]. The residue sequence is further decided with ProteinMPNN[29], and the side-chain conformation is added with Rosetta[99] side-chain packing.

**RFDiffusionAA** [16] ^4^ is the latest version of RFDiffusion which combines a residue-based representation of amino acids and atomic representations of all other groups to model protein-small molecules/metals/nucleic acids/covalent modification complexes. Starting from random distributions of amino acid residues surrounding target small molecules, RFDiffusionAA can directly generate the small molecule binding protein backbone. Furthermore, with LigandMPNN [30], the latest version of ProteinMPNN[29], we can assign residue types and predict sidechain conformations considering the protein-ligand interactions. Experiments in RFDiffusionAA [16] show that the generated protein by RFDiffusionAA has better binding affinity than those obtained by RFDiffusion with auxiliary potential.

**dyMEAN** [25] ^5^ is an end-to-end full-atom model for E(3)-equivariant antibody design given the epitope and the incomplete sequence of the antibody. Its previous version, MEAN [34], only considers the backbone atoms, while dyMEAN considers the complete atom structure and performs better on downstream tasks. Generally, dyMEAN co-designs antibody sequence and structure via a multi-round progressive full-shot refinement manner, which is more efficient than auto-regressive or diffusion-based approaches. An adaptive multi-channel equivariant encoder is used in dyMEAN, which can process protein residues of variable sizes when considering full atoms. To adapt dyMEAN to our pocket design task, we replace the antigen with the target ligand molecule to provide the context information for pocket generation.

**FAIR** [24] ^6^ is our previous method for full atom pocket sequence-structure co-design. FAIR operates in two steps, proceeding in a coarse-to-fine manner (backbone refinement to full atoms refinement, including side chains) for full-atom generation. In FAIR, residue types and atom coordinates are updated using a hierarchical graph transformer composed of a residue-level and atom-level encoder.

#### Proof of E(3)-Equivariance of PocketGen

The E(3)-transformation on the euclidean coordinate ***x*** ∈ ℝ^3^ can be represented as: *T*_*g*_ · ***x*** = ***Ox***+***t***, where ***O*** ∈ ℝ^3^ is the orthogonal transformation matrix, ***t*** ∈ R^3^ is the translation vector. Implementing *T*_*g*_ on a coordinate matrix ***X*** ∈ ℝ^*n*×3^ means transforming each coordinate (i.e., each row) with *T*_*g*_. PocketGen has the desirable property of **E(3)-equivariance** as follow:

##### Theorem 1.

*Denote the E(3)-transformation as T*_*g*_ *and the generative process of PocketGen as*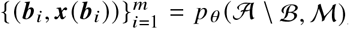, *where* 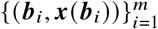 *indicates the designed protein pocket seqeuce and structure. We have* {(***b***_*i*_, *T*_*g*_ · ***x*** (***b***_*i*_))}^*m*^ = 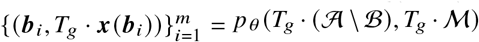. *Here, T*_*g*_ · (𝒜 \ ℬ) *and T*_*g*_· ℳ *denote applying E(3)-transformation on the protein structures except for the pocket region and the molecular structure respectively*.

Then we prove the E(3)-equivariance of each module in PocketGen as follows.

##### Lemma 1.

*Denote the bilevel attention module as* 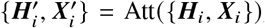, *then it is E(3)-equivariant. Namely, for any E(3)-transformation T*_*g*_, *we have* 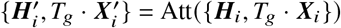.

*Proof*. The key to the proof of Lemma 1 is to prove that the propagation in Eq. 2-9 is E(3)-invariant on ***H***_*i*_ and E(3)-equivariant on ***X***_*i*_. The correlation ***R***_*i j*_ between block *i* and block *j* in Eq. 3 is E(3)-invariant because all the inputs, including the query ***Q***_*i*_, the key ***K*** _*j*_, and the distance matrices ***D***_*i j*_, are not influenced by the geometric transformation *T*_*g*_. Therefore, we can immediately derive that the atom-level attention ***α***_*i j*_ in Eq. 4 is E(3)-invariant. Similarly, the residue/ligand-level attention *β*_*i j*_ in Eq. 6 is E(3)-invariant because it only operates on *r*_*i j*_ in Eq. 5 which aggregates ***α***_*i j*_. Finally, we can derive the E(3)-invariance on ***H*** and the E(3)-equivariance on ***X*** (ligand or pocket residues) as below:

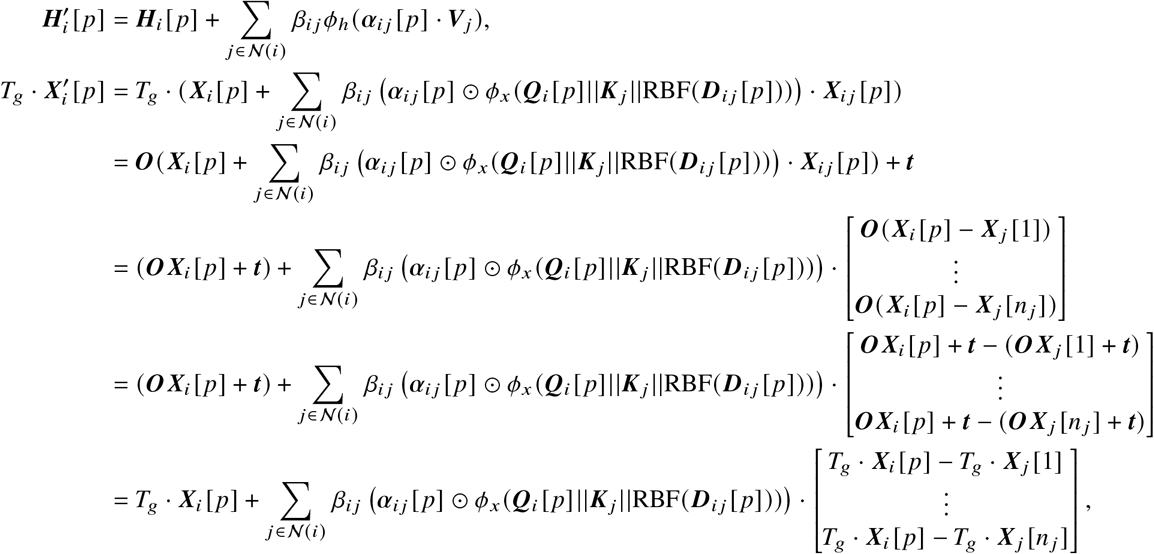

which concludes the proof of Lemma 1.

##### Lemma 2.

*Denote the equivariant feed-forward network as as* 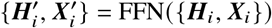, *then it is E(3)-equivariant. Namely, for any E(3)-transformation T*_*g*_, *we have*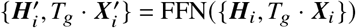.

*Proof*. The proof of Lemma 2 focuses on the single-atom updates in Eq. 10-13. First, it is easy to obtain the E(3)-equivariance of the centroid in Eq. 10:

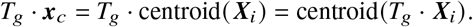

Then we have the following equation on the relative coordinate Δ***x*** in Eq. 11:

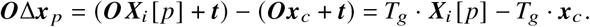

Then we can obtain the E(3)-invariance of ***r*** _*p*_ in Eq. 11:

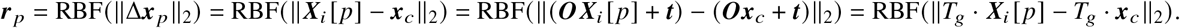

Finally we can derive the E(3)-invariance on ***H***^′^ [ *p*] and the E(3)-equivariance on 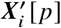:

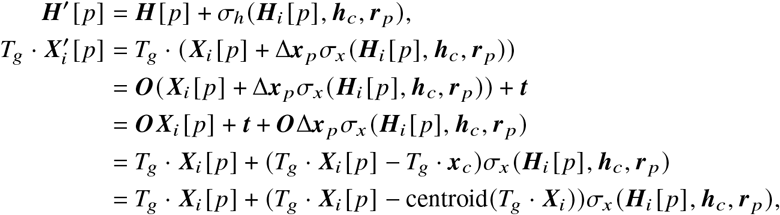

which concludes the proof of Lemma 2

The sequence refinement module with the pretrained protein language models and adapters operates on the scaler residue representations unaffected by the geometric transformation *T*_*g*_. To sum up, with Lemma 1-2 at hand, it is obvious to deduce the E(3)-equivariance of the PocketGen.

## Supplementary Figures

**Figure S1.**
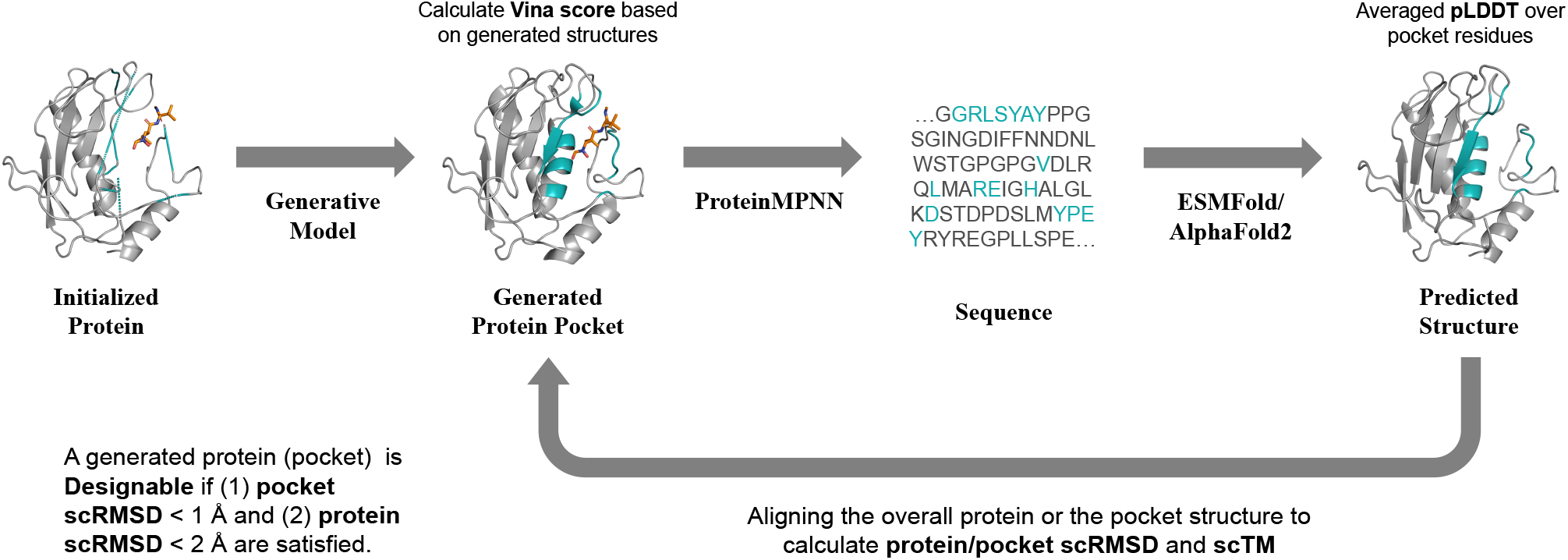
Schematic of computing metrics with respect to the generated pocket structures. black parts refer to the pocket, while the others are gray. The generative model refers to PocketGen or baseline methods leveraged to generate the protein pocket. The Vina score is calculated based on the generated protein-ligand complex. To calculate self-consistency scores, we first use ProteinMPNN to derive the residue sequence and then use ESMFold/AlphaFold2 to predict the structure. By aligning the predicted structure with the generated structure, we can obtain the protein/pocket scRMSD. Similarly, we can obtain the scTM score of the protein structure. We also report the averaged pLDDT of the pocket residues from ESMFold.

**Figure S2.**
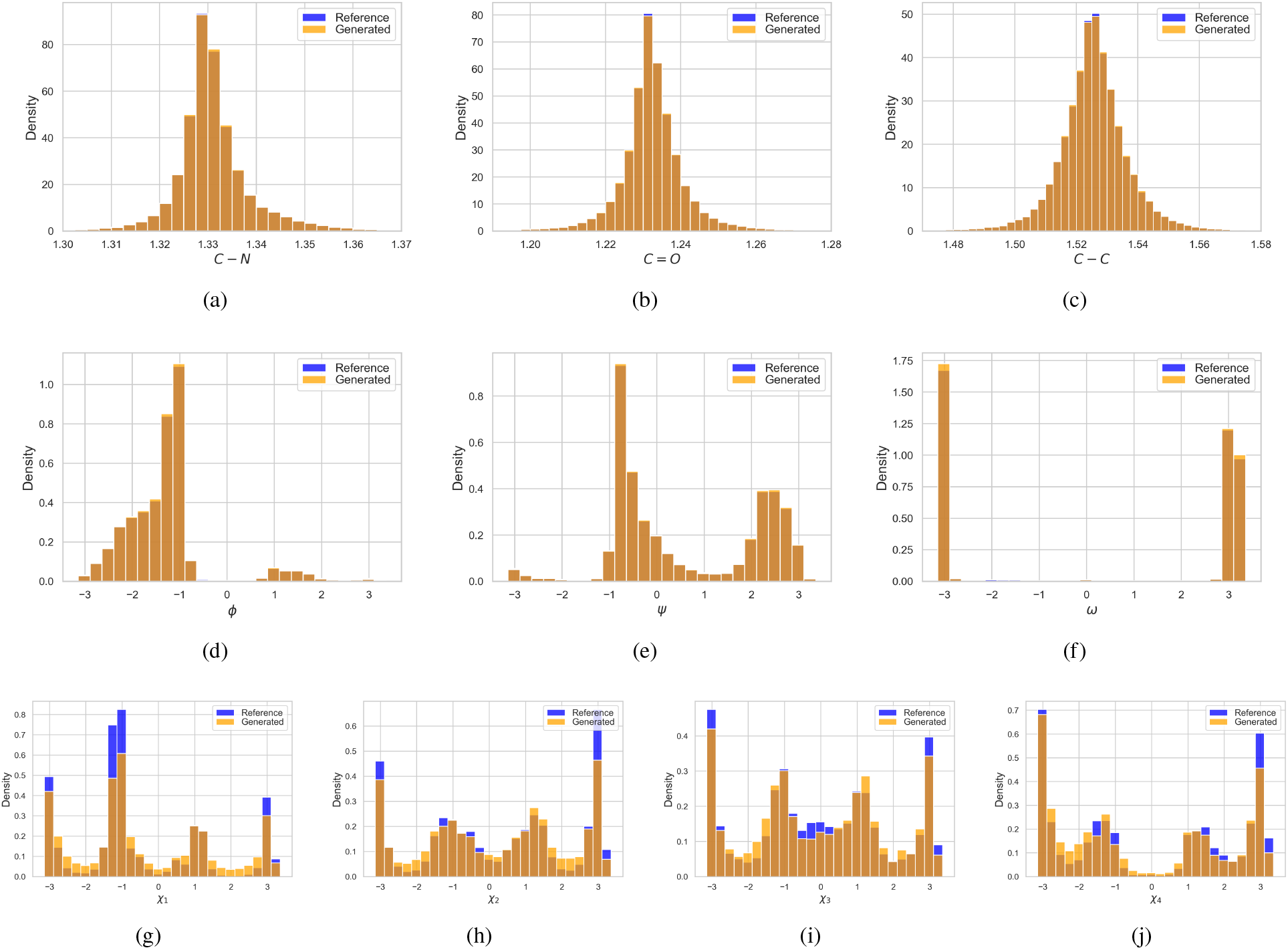
The bond length and dihedral angle distributions of the generated pockets and the reference (training set of CrossDocked dataset).

**Figure S3.**
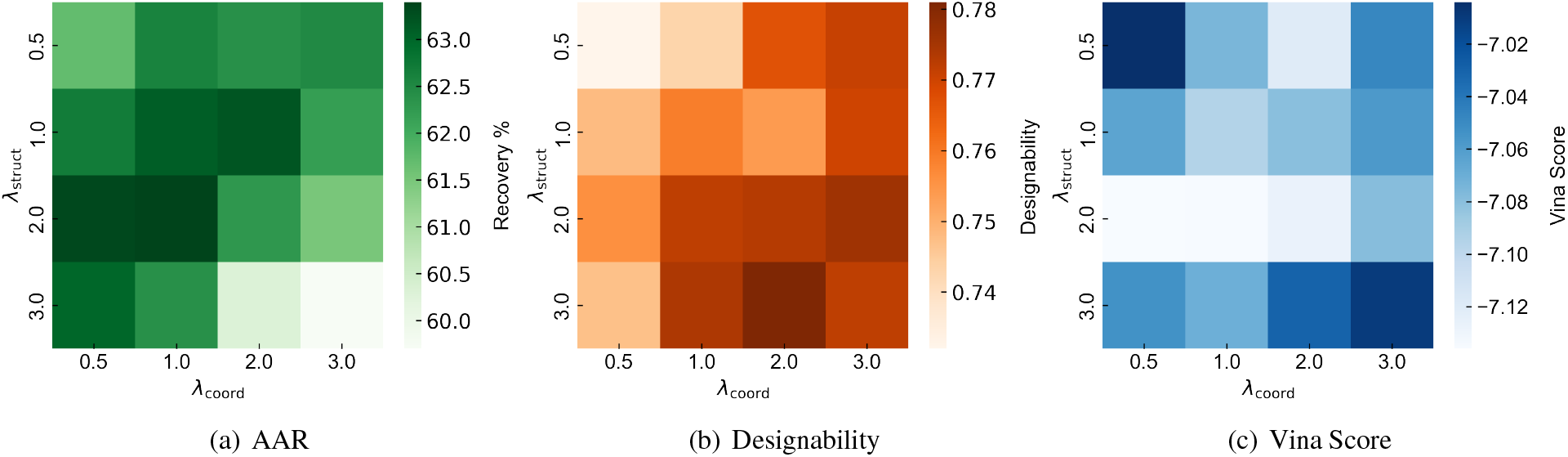
Hyperparameter Analysis. Influence of hyperparameters (loss weights ℒ _coord_ and ℒ _struct_) on PocketGen’s performance (AAR, Designability, and Vina Score) on the CrossDocked dataset. We perform a grid search over {0.5, 1.0, 2.0, 3.0} and choose these hyperparameters based on the validation performance to select the specific parameter values. In the default setting, we set *λ*_coord_ to 1.0 and *λ*_struct_ to 2.0.

## Supplementary Tables

**Table S1.** Benchmarking PocketGen and other approaches for pocket generation on two datasets. Reported are average and standard deviation values across three independent runs (random seeds on the same dataset). The best results are bolded.

| Model | CrossDocked |  |  | Binding MOAD |  |  |
| --- | --- | --- | --- | --- | --- | --- |
|  | AAR (↑) | Designability (↑) | Vina (↓) | AAR (↑) | Designability (↑) | Vina (↓) |
| Test set | - | 0.77 | -7.016 | - | 0.79 | -8.076 |
| DEPACT | 31.52±3.26% | 0.68±0.04 | -6.632±0.18 | 35.30±2.19% | 0.67±0.06 | -7.571±0.15 |
| dyMEAN | 38.71±2.16% | 0.71±0.03 | -6.855±0.06 | 41.22±1.40% | 0.70±0.03 | -7.675±0.09 |
| FAIR | 40.16±1.17% | 0.73±0.02 | -7.015±0.12 | 43.68±0.92% | 0.72±0.05 | -7.930±0.15 |
| RFDiffusion | 46.57±2.07% | 0.74±0.01 | -6.936±0.07 | 45.31±2.73% | 0.75±0.05 | -7.942±0.14 |
| RFDiffusionAA | 50.85±1.85% | 0.75±0.03 | -7.012±0.09 | 49.09±2.49% | 0.78±0.03 | -8.020±0.11 |
| PocketGen | **63.40±1.64%** | **0.77±0.02** | **-7.135±0.08** | **64.43±2.35%** | **0.80±0.04** | **-8.112±0.14** |

**Table S2.** The top 1/3/5/10 generated protein pocket (ranked by Vina score) with designability on the CrossDocked dataset. The success rate measures the percentage of protein that the model can generate pockets with higher affinity than the reference ones in the datasets. We report the average plDDT of the predicted pocket, the scRMSD of the pocket backbone coordinates, and the change of the scTM score of the whole protein. ESMFold means the scores are calculated with AlphaFold2 as the folding tool. The plDDT, scRMSD, and ΔscTM for PocketOpt are not reported, as PocketOpt keeps protein backbone structures fixed. We 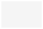 use to mark the results of affinity-related metrics, 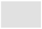 for pocket-structure related metrics, and 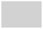 for whole protein structure metrics. We report the means and standard deviations over three independent runs with random seeds. The best results are indicated in **bold**.

|  | DEPACT | dyMEAN | FAIR | RFDiffusion | RFAA | PocketGen |
| --- | --- | --- | --- | --- | --- | --- |
| Top-1 generated protein pocket |  |  |  |  |  |  |
| pLDDT (ESMFold) (↑) | 82.130±0.187 | 83.327±0.206 | 83.254±0.412 | 84.559±0.230 | 86.346±0.238 | **87.065**±0.180 |
| scRMSD (ESMFold) (↓) | 0.705±0.023 | 0.703±0.024 | 0.680±0.019 | 0.676±0.017 | 0.654±0.011 | **0.642**±0.008 |
| ΔscTM (ESMFold) (↑) | -0.009±0.002 | -0.004±0.003 | -0.011±0.004 | 0.014±0.005 | 0.021±0.003 | **0.030**±0.004 |
| ΔscTM (ESMFold+co) (↑) | -0.015±0.005 | -0.016±0.003 | -0.023±0.005 | - | - | **0.014**±0.003 |
| Top-3 generated protein pockets |  |  |  |  |  |  |
| pLDDT (ESMFold) (↑) | 81.991±0.303 | 82.825±0.218 | 83.145±0.297 | 84.636±0.186 | 86.224±0.190 | **86.450**±0.075 |
| scRMSD (ESMFold) (↓) | 0.706±0.025 | 0.724±0.024 | 0.685±0.017 | 0.679±0.014 | **0.653**±0.012 | 0.655±0.007 |
| ΔscTM (ESMFold) (↑) | -0.014±0.003 | -0.010±0.004 | -0.013±0.005 | 0.019 ±0.004 | 0.020±0.003 | **0.024** ±0.001 |
| ΔscTM (ESMFold+co) (↑) | -0.016±0.003 | -0.022±0.002 | -0.026±0.003 | - | - | **0.012**±0.002 |
| Top-5 generated protein pockets |  |  |  |  |  |  |
| pLDDT (ESMFold) (↑) | 82.070±0.276 | 82.910±0.231 | 83.168±0.208 | 84.320±0.219 | 85.735±0.087 | **86.414**±0.110 |
| scRMSD (ESMFold) (↓) | 0.717±0.019 | 0.725±0.014 | 0.690±0.012 | 0.680±0.012 | 0.656±0.006 | **0.654**±0.010 |
| ΔscTM (ESMFold) (↑) | -0.018±0.004 | -0.007 ±0.002 | -0.024±0.002 | 0.016±0.001 | 0.019±0.001 | **0.027**±0.001 |
| ΔscTM (ESMFold+co) (↑) | -0.014±0.004 | -0.023±0.003 | -0.027±0.002 | - | - | **0.015**±0.002 |
| Top-10 generated protein pockets |  |  |  |  |  |  |
| pLDDT (ESMFold) (↑) | 81.580±0.232 | 82.771±0.203 | 83.048±0.165 | 84.234±0.195 | 85.377±0.142 | **86.090**±0.124 |
| scRMSD (ESMFold) (↓) | 0.710±0.014 | 0.734±0.013 | 0.705±0.013 | 0.684±0.012 | 0.672±0.007 | **0.657**±0.005 |
| ΔscTM (ESMFold) (↑) | -0.025±0.002 | -0.014±0.001 | -0.026±0.002 | 0.014±0.003 | 0.019±0.000 | **0.022**±0.001 |
| ΔscTM (ESMFold+co) (↑) | -0.015±0.003 | -0.022±0.002 | -0.026±0.003 | - | - | **0.013**±0.001 |

**Table S3.** The top 1/3/5/10 generated protein pocket (ranked by Vina score) with designability on the Binding MOAD dataset. Besides Vina score, we also calculate MM-GBSA to evaluate the binding affinity. The success rate measures the percentage of protein that the model can generate pockets with higher affinity than the reference ones in the datasets. We report the average plDDT of the predicted pocket, the scRMSD of the pocket backbone coordinates, and the change of the scTM score of the whole protein. co indicates codesign, where codesign methods directly use the designed sequence for consistency calculation. The plDDT, scRMSD, and ΔscTM for PocketOpt are not reported, as PocketOpt keeps protein backbone structures fixed. We use 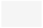 to mark the results of affinity-related metrics,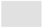 for pocket-structure related metrics, and 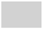 for whole protein structure metrics. We report the means and standard deviations over three independent runs with random seeds. The best results are indicated in **bold**.

| Methods: | PocketOpt | DEPACT | dyMEAN | FAIR | RFDiffusion | RFAA | PocketGen |
| --- | --- | --- | --- | --- | --- | --- | --- |
| Top-1 generated protein pocket |  |  |  |  |  |  |  |
| Vina score (↓) | -9.828±0.120 | -9.216±0.071 | -9.352±0.082 | -9.530±0.064 | -9.901±0.055 | -10.120±0.063 | **-10.322±0.057** |
| MM-GBSA (↓) | -62.293±0.819 | -55.180±0.906 | -55.504±0.628 | -59.851±0.723 | -63.440±0.836 | -64.305±1.04 | **-67.862±0.618** |
| Success Rate (↑) | 0.891±0.024 | 0.784±0.045 | 0.779±0.02 | 0.841±0.033 | 0.905±0.030 | 0.923±0.027 | **0.952±0.016** |
| pLDDT (ESMFold) (↑) | - | 83.145±0.426 | 83.280±0.247 | 83.536±0.217 | 85.068±0.335 | **87.232±0.314** | 87.106±0.224 |
| scRMSD (ESMFold) (↓) | - | 0.642±0.024 | 0.622±0.017 | 0.620±0.025 | 0.581±0.018 | **0.572±0.016** | 0.575±0.013 |
| ΔscTM (ESMFold) (↑) | - | -0.004±0.001 | -0.001±0.002 | 0.008±0.003 | 0.025±0.005 | **0.031±0.005** | 0.030±0.004 |
| ΔscTM (ESMFold+co) (↑) | - | -0.025±0.003 | -0.019±0.004 | -0.008±0.005 | - | - | 0.004±0.003 |
| Top-3 generated protein pocket |  |  |  |  |  |  |  |
| Vina score (↓) | -9.403±0.102 | -8.876±0.064 | -8.971±0.076 | -9.234±0.050 | -9.589±0.048 | -9.634±0.045 | **-10.135±0.039** |
| MM-GBSA (↓) | -58.204±0.739 | -51.255±0.660 | -52.371±0.540 | -54.946±0.632 | -57.235±0.803 | -60.801±0.829 | **-64.108±0.421** |
| pLDDT (↑) | - | 82.769±0.370 | 82.131±0.263 | 82.826±0.207 | 84.912±0.328 | 85.920±0.301 | **86.535±0.220** |
| scRMSD (ESMFold) (↓) | - | 0.645±0.023 | 0.620±0.015 | 0.596±0.014 | 0.580±0.016 | 0.574±0.015 | **0.573±0.011** |
| ΔscTM (ESMFold) (↑) | - | -0.009±0.002 | -0.004±0.003 | 0.004±0.002 | 0.020±0.005 | 0.025±0.004 | **0.029±0.003** |
| ΔscTM (ESMFold+co) (↑) | - | -0.027±0.002 | -0.020±0.003 | -0.007±0.002 | - | - | **0.005±0.002** |
| Top-5 generated protein pocket |  |  |  |  |  |  |  |
| Vina score (↓) | -9.260±0.091 | -8.759±0.050 | -8.842±0.050 | -9.195±0.043 | -9.478±0.045 | -9.569±0.035 | **-9.950±0.030** |
| MM-GBSA (↓) | -55.728±0.536 | -49.204±0.745 | -50.289±0.431 | -52.152±0.577 | -55.490±0.830 | -56.306±0.687 | **-62.970±0.412** |
| pLDDT (ESMFold) (↑) | - | 81.939±0.261 | 81.915±0.163 | 82.346±0.219 | 85.150±0.322 | 85.830±0.205 | **86.438±0.160** |
| scRMSD (ESMFold) (↓) | - | 0.639±0.015 | 0.626±0.014 | 0.612±0.015 | 0.587±0.010 | 0.583±0.011 | **0.581±0.008** |
| ΔscTM (ESMFold) (↑) | - | -0.007±0.001 | -0.002±0.002 | 0.003±0.002 | 0.024±0.003 | 0.024±0.003 | **0.026±0.001** |
| ΔscTM (ESMFold+co) (↑) | - | -0.027±0.004 | -0.023±0.001 | -0.005±0.002 | - | - | **0.005±0.002** |
| Top-10 generated protein pocket |  |  |  |  |  |  |  |
| Vina score (↓) | -8.981±0.087 | -8.632±0.056 | -8.690±0.051 | -8.827±0.063 | -9.268±0.042 | -9.320±0.054 | **-9.630±0.037** |
| MM-GBSA (↓) | -51.337±0.505 | -45.480±0.519 | -47.804±0.420 | -48.203±0.625 | -53.660±0.522 | -55.763±0.643 | **-60.106±0.284** |
| pLDDT (ESMFold) (↑) | - | 81.632±0.270 | 81.774±0.264 | 82.739±0.340 | 84.582±0.191 | 85.337±0.229 | **86.456±0.079** |
| scRMSD (ESMFold) (↓) | - | 0.644±0.012 | 0.626±0.014 | 0.592±0.013 | 0.589±0.014 | 0.586±0.007 | **0.584±0.005** |
| ΔscTM (ESMFold) (↑) | - | -0.003±0.002 | -0.006±0.001 | 0.006±0.003 | 0.012±0.002 | 0.018±0.002 | **0.027±0.001** |
| ΔscTM (ESMFold+co) (↑) | - | -0.029±0.002 | -0.021±0.001 | -0.008±0.002 | - | - | **0.006±0.003** |

**Table S4.**
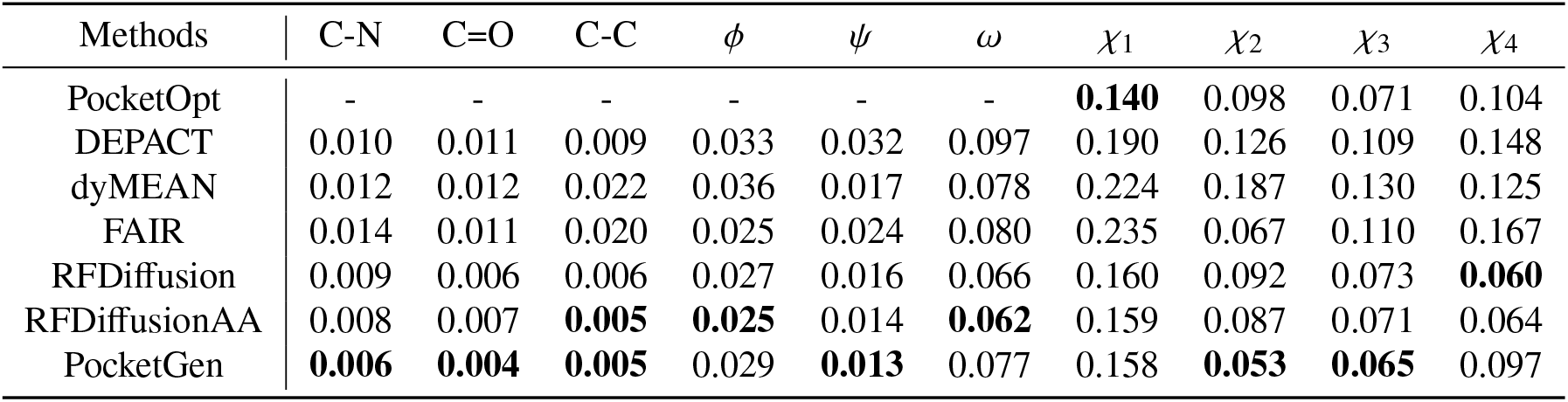
Substructure analysis of the generated molecules. We consider three covalent bonds in the backbone (C-N, C=O, and C-C), three conventional dihedral angles in the backbone (*ϕ, ψ, ω* [58]), and four dihedral angles in the side chains (*χ*_1_, *χ*_2_, *χ*_3_, *χ*_4_ [59]). The KL divergence of the bond lengths and dihedral angles between the training set (CrossDocked) and the generated pockets are calculated following previous works[60, 61]. Since PocketOpt sets the pocket backbone fixed, the corresponding backbone metrics are not calculated. The best results are indicated in **bold**.

**Table S5.** **Ablations studies of PocketGen. PocketGen w/o ligand update** indicates the ligand structure is fixed during the generation of pockets by PocketGen. **PocketGen w/o residue-level attention** denotes only the basic atom-level attention performed in the bilevel attention module. **PocketGen w/o pLM** means no pretrained protein language model is leveraged for sequence refinement. Moreover, **PocketGen w/ EGNN/GVP/GMN** uses EGNN[67], GVP[68], and GMN[69] respectively to replace the bilevel graph transformer in PocketGen as the structural encoder for comparison. We report the means and standard deviations over three different runs (%). The best results are bolded.

| Model | CrossDocked |  |  | Binding MOAD |  |  |
| --- | --- | --- | --- | --- | --- | --- |
| | AAR (↑) | Designability (↑) | Vina (↓) | AAR (↑) | Designability (↑) | Vina (↓) |
| PocketGen w/o ligand update | 59.20±1.56% | 0.75±0.03 | -6.838±0.10 | 59.61±2.13% | 0.77±0.05 | -7.865±0.12 |
| PocketGen w/o residue-level attention | 58.19±1.78% | 0.66±0.04 | -6.924±0.08 | 57.30±2.28% | 0.67±0.04 | -7.805±0.10 |
| PocketGen w/o pLM | 42.70±1.45% | 0.71±0.05 | -6.894±0.07 | 44.23±1.31% | 0.69±0.06 | -7.787±0.12 |
| PocketGen w/ EGNN encoder | 58.85±1.21% | 0.71±0.02 | -6.916±0.03 | 58.64±1.14% | 0.72±0.06 | -7.804±0.11 |
| PocketGen w/ GVP encoder | 61.10±0.90% | 0.72±0.04 | -6.930±0.08 | 59.97±1.40% | 0.75±0.05 | -7.883±0.10 |
| PocketGen w/ GMN encoder | 60.49±1.33% | 0.68±0.03 | -6.947±0.05 | 59.82±1.67% | 0.74±0.04 | -7.869±0.08 |
| PocketGen | **63.40±1.64%** | **0.77±0.02** | **-7.135±0.08** | **64.43±2.35%** | **0.80±0.04** | **-8.112±0.14** |

**Table S6.**
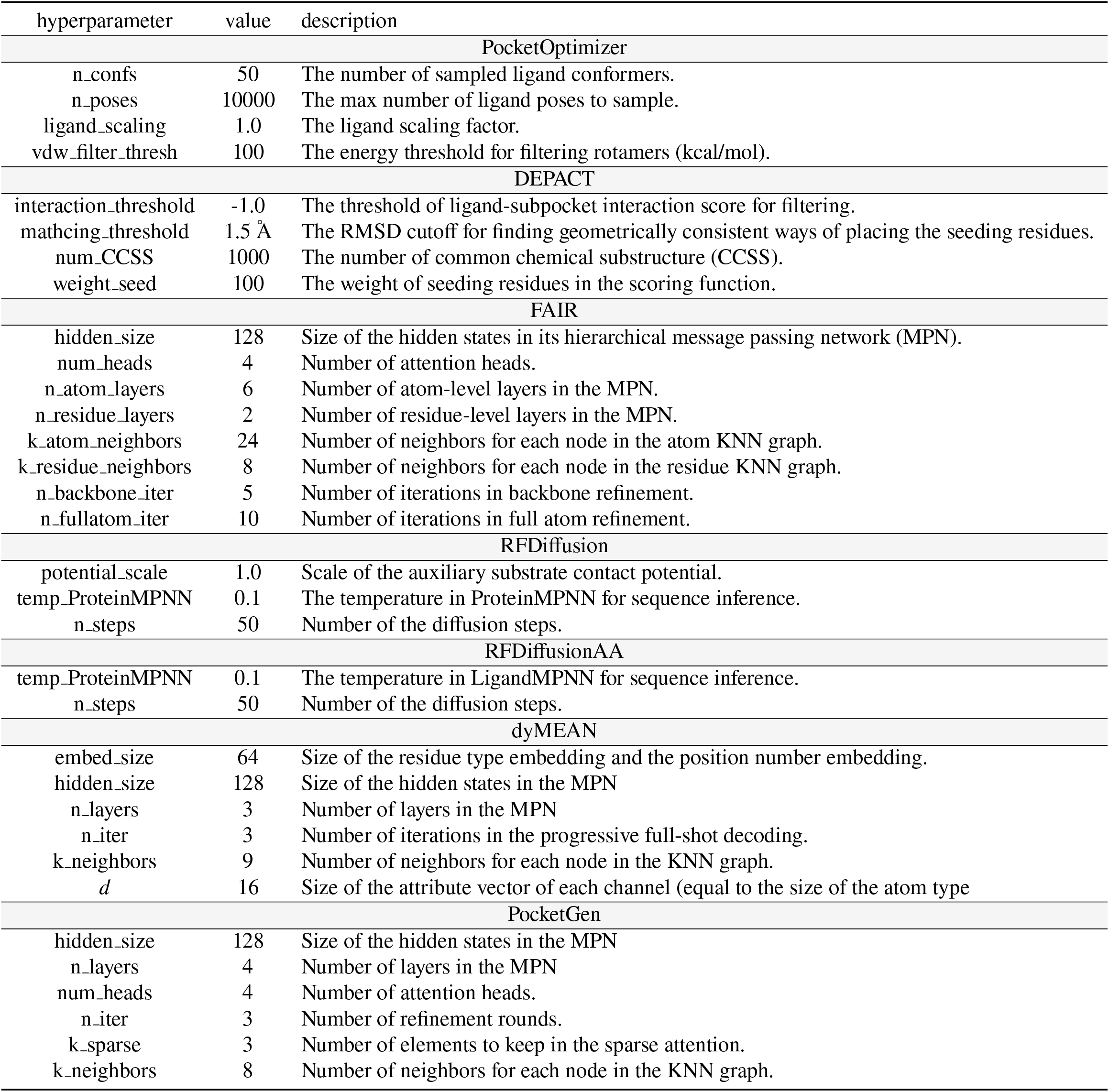
Hyperparameters for the baselines and our PocketGen.

| hyperparameter | value | description |
| --- | --- | --- |
| PocketOptimizer |  |  |
| n_confs | 50 | The number of sampled ligand conformers. |
| n_poses | 10000 | The max number of ligand poses to sample. |
| ligand_scaling | 1.0 | The ligand scaling factor. |
| vdw_filter_thresh | 100 | The energy threshold for filtering rotamers (kcal/mol). |
| DEPACT |  |  |
| interaction_threshold | -1.0 | The threshold of ligand-subpocket interaction score for filtering. |
| mathcing_threshold | 1.5 Å | The RMSD cutoff for finding geometrically consistent ways of placing the seeding residues. |
| num_CCSS | 1000 | The number of common chemical substructure (CCSS). |
| weight_seed | 100 | The weight of seeding residues in the scoring function. |
| FAIR |  |  |
| hidden_size | 128 | Size of the hidden states in its hierarchical message passing network (MPN). |
| num_heads | 4 | Number of attention heads. |
| n_atom_layers | 6 | Number of atom-level layers in the MPN. |
| n_residue_layers | 2 | Number of residue-level layers in the MPN. |
| k_atom_neighbors | 24 | Number of neighbors for each node in the atom KNN graph. |
| k_residue_neighbors | 8 | Number of neighbors for each node in the residue KNN graph. |
| n_backbone_iter | 5 | Number of iterations in backbone refinement. |
| n_fullatom_iter | 10 | Number of iterations in full atom refinement. |
| RFDiffusion |  |  |
| potential_scale | 1.0 | Scale of the auxiliary substrate contact potential. |
| temp_ProteinMPNN | 0.1 | The temperature in ProteinMPNN for sequence inference. |
| n_steps | 50 | Number of the diffusion steps. |
| RFDiffusionAA |  |  |
| temp_ProteinMPNN | 0.1 | The temperature in LigandMPNN for sequence inference. |
| n_steps | 50 | Number of the diffusion steps. |
| dyMEAN |  |  |
| embed_size | 64 | Size of the residue type embedding and the position number embedding. |
| hidden_size | 128 | Size of the hidden states in the MPN |
| n_layers | 3 | Number of layers in the MPN |
| n_iter | 3 | Number of iterations in the progressive full-shot decoding. |
| k_neighbors | 9 | Number of neighbors for each node in the KNN graph. |
| d | 16 | Size of the attribute vector of each channel (equal to the size of the atom type |
| PocketGen |  |  |
| hidden_size | 128 | Size of the hidden states in the MPN |
| n_layers | 4 | Number of layers in the MPN |
| num_heads | 4 | Number of attention heads. |
| n_iter | 3 | Number of refinement rounds. |
| k_sparse | 3 | Number of elements to keep in the sparse attention. |
| k_neighbors | 8 | Number of neighbors for each node in the KNN graph. |

## Footnotes

1 https://github.com/Hoecker-Lab/pocketoptimizer

2 https://github.com/chenyaoxi/DEPACT_PACMatch

3 https://github.com/RosettaCommons/RFdiffusion

4 https://github.com/baker-laboratory/rf_diffusion_all_atom

5 https://github.com/THUNLP-MT/dyMEAN

6 https://github.com/zaixizhang/FAIR

